# Two germ granule eIF4E isoforms reside in different mRNPs to hand off *C elegans* mRNAs from translational repression to activation

**DOI:** 10.1101/2024.05.24.595216

**Authors:** Gita Gajjar, Hayden P. Huggins, Eun Suk Kim, Weihua Huang, Frederic X. Bonnet, Dustin L. Updike, Brett D. Keiper

## Abstract

We studied the function of translation factor eIF4E isoforms in regulating mRNAs in germ cell granules/condensates. Translational control of mRNAs plays an essential role in germ cell gene regulation. Messenger ribonucleoprotein (mRNP) complexes assemble on mRNAs as they move from the nucleus into perinuclear germ granules to exert both positive and negative post-transcriptional regulation in the cytoplasm. In *C. elegans*, germ granules are surprisingly dynamic mRNP condensates that remodel during development. Two eIF4E isoforms (called IFE-1 and IFE-3), eIF4E-Interacting Proteins (4EIPs), RBPs, DEAD-box helicases, polyadenylated mRNAs, Argonautes and miRNAs all occupy positions in germ granules. Affinity purification of IFE-1 and IFE-3 allowed mass spectrometry and mRNA-Seq to identify the proteins and mRNAs that populate stable eIF4E mRNPs. We find translationally controlled mRNAs (e.g. *pos-1, mex-3, spn-4,* etc.) enriched in IFE-3 mRNPs, but excluded from IFE-1 mRNPs. These mRNAs also require IFE-1 for efficient translation. The findings support a model in which oocytes and embryos utilize the two eIF4Es for opposite purposes on critically regulated germline mRNAs. Careful colocalization of the eIF4Es with other germ granule components suggests an architecture in which GLH-1, PGL-1 and the IFEs are stratified to facilitate sequential interactions for mRNAs. Biochemical characterization demonstrates opposing yet cooperative roles for IFE-3 and IFE-1 to hand-off of translationally controlled mRNAs from the repressed to the activated state, respectively. The model involves eIF4E mRNPs shuttling mRNAs through nuclear pore-associated granules/condensates to cytoplasmic ribosomes.

## Introduction

Protein synthesis is regulated both qualitatively and quantitatively in nearly all dynamic biological settings. This macromolecular process requires a series of coordinated interactions of RNA and protein, including the ribosome itself and the ribonucleoprotein complexes (mRNPs) forming on mRNAs (1–8). Interactions within these RNA-protein complexes are remodeled substantially during translation initiation because mRNAs that have just finished splicing emerge from the nucleus as complex mRNPs. Multiple eukaryotic initiation factors (eIFs) assemble on the both the small ribosomal subunit (40S) and the substrate mRNP. A ternary complex composed of eukaryotic initiation factor 2 (eIF2) binding to Met-tRNAi and GTP binds to free 40S, assisted by eIF1, eIF1A, eIF3, and eIF5, to make a 43S pre-initiation complex (43S PIC) (2,8). Meanwhile, eIF4F factors (eIF4E, eIF4G, eIF4A, eIF4B) typically assemble on the 5’ m^7^GpppN mRNA cap and associate via eIF3 with the 43S PIC to make a committed 48S PIC that scans 5’ to 3’ toward the initiation codon (8). Upon recognition of AUG by eIF2/ternary complex, eIF5B, and ABCE1 promote 60S ribosomal subunit joining to form an 80S monosome that is competent to begin synthesis (elongation; (9,10). Control of mRNA translation initiation generally occurs at the eIF2- and/or eIF4-catalyzed steps. Regulation of cap-dependent (eIF4) recruitment, involving eIF4E-interacting proteins (4EIPs), mTOR kinase, and eIF4G isoform selection, is a highly-coordinated mode of gene expression that has implications for plant and animal development, reproduction (3,4,11,12), oncogenic transformation (13–15), and neuronal functions (16–18). Besides translation initiation, a few eIFs take part in other aspects of mRNA and small RNA (miRNA, piRNA) life. Isoforms of the cap-binding protein eIF4E, the helicase eIF4A, and eIF3 subunits have additional non-canonical roles that exert repression, destabilization, or mechanism shift (e.g. cap-independent translation of viral/apoptotic mRNAs), as well as feedback from the protein synthetic apparatus to trigger physiological or developmental cell fate changes (3,5,12,19–29).

There are many unique RNA regulatory mechanisms at work than in germ cells, the immortal lineage of precursors to reproduction. Germ cells use extensive mRNA translational control to drive both their temporal maintenance of the undifferentiated state and their differentiation during gametogenesis and eventual embryogenesis. mRNA translational control is mediated by RNA-binding proteins, miRNAs, translation initiation factors, and interacting proteins in mRNP structures (12,30). This activity begins as mRNAs emerge from the nucleus through the nuclear pore complex and into perinuclear germ granules on their way to the cytoplasmic ribosomes. Germ granules are dense, membrane-less, self-driven condensates composed largely of RNA and protein that exist in the germ lineage of all known sexual reproducing species (31–33). These condensates physically demix from their soluble context, becoming a pseudo-stable separated phase that is simultaneously very dynamic. Physical properties in the condensate cause their cargo macromolecules to undergo smooth phase transitions and become structured and organized (33,34). In *C. elegans*, germ granule condensates directly overlay nuclear pore complexes (NPCs). They receive exported mRNAs, micro RNAs (miRNA) and piwi RNAs (piRNA) to organize, localize, enzymatically manipulate, and decorate them with RBPs in mRNP form. Within germ granules themselves there is now evidence of three or more subcompartments: Z granules, Mutator foci, P-body-like granules and SIMR foci, each with potential importance to mRNA translational control or turnover (33,35,36). Proteins with high residency time in the germ granule provide the structure on which the mRNPs assemble and usually encode intrinsically disordered regions, low complexity domains, RNA binding, and repetitive elements (32,37,38). These traits lend themselves to RNA-protein remodeling and hand off of cargo that we propose move mRNAs through the periplasmic condensate toward ribosomes. The remodeling serves to license the mRNP for one of three outcomes: translational repression (implicit stabilization), translational activation, or mRNA turnover. It is notable that relative transcription quiescence in germ cells necessitates that maternal mRNAs are expressed long before their protein products are needed for subsequent gametogenesis and the oocyte/embryo transition (OET). Their temporal synthesis is governed by well-studied mRNA translational regulation events (24,30,39) in which new roles for mRNA condensate residence often plays a role.

There are five isoforms of eIF4E expressed in the nematode *C. elegans* (IFEs −1, −2, −3, −4 and −5) that share strong sequence homology but have largely non-overlapping functions (12,29). Our earlier studies identified small subpopulations of mRNAs that are uniquely translationally controlled by each of four different IFE isoforms (40–44). Two of these eIF4Es (IFE-1 and IFE-3) are enriched in germ granules and can exert both positive and negative translational regulation on their preferred mRNA targets (12,24). The current study extends the characterization of IFE-1 and IFE-3 to the mRNPs in which they reside and the mRNAs that they regulate. Here we identify the proteins and mRNAs that uniquely populate two distinct eIF4E mRNPs. The identities of the proteins in each mRNP point to multiple cap-binding roles, not only for mRNA translational control, but also for small RNA processing and mRNA turnover pathways. We further demonstrate that IFE-1 and IFE-3 along with their unique 4EIPs have opposing functional roles in the translational control of cargo mRNAs. Comparison of mRNA populations in each stable IFE mRNP to mRNAs previous identified to undergo IFE-mediated translational activation or repression also gives evidence for a hand-off of germ cell mRNAs between the two germ granule eIF4Es on the way to recruitment to ribosomes. We propose that translational control by eIF4E mRNPs preselects the active and repressive translation fate of mRNAs before they emerge from germ granule complexes and facilitates rapid changes in mRNA utilization or decay in response to developmental stimuli.

## Methods

### Strains and maintenance

C. elegans Bristol var. N2 was used as the wild type strain in all experiments unless otherwise noted. Strains used in this study and their genotypes are described in Table 1. Strains expressing fluorescent CRISPR/Cas9 fusions to the pos-1, mex-3, spn-4 and lin-41 (DG4222, DG4269, DG4739) were generous provided by Dr. David Greenstein (University of Minnesota). Other strains were obtained from the Caenorhabditis Genetics Center (CGC, University of Minnesota) which is funded by NIH Office of Research Infrastructure Programs [grant number P40 OD010440]. Strains containing CRISPR/Cas9 fusions of ife-1, ife-3 an/or other germ granule protein genes, as well as null deletions in ife-1 and ife-3 were described in detail in (40) and (45). These deletion strains are differentially temperature sensitive. Hermaphrodites homozygous for the ife-1(bn127) deletion were sterile at 25C, but could produce viable embryos at 20C with markedly reduced fertility (46). Worms bearing the ife-3(ok191) null mutation exhibit embryonic lethality and a few hermaphrodite escapers with masculinized gonads (40). To overcome this, ife-3(ok191) was balanced with eT1 containing myo-2::gfp and the strains were maintained as heterozygotes. Homozygous ife-3 null worms were identified as lacking GFP in the pharynx and were used for relevant experiments in this study as described (40). Newly crossed strains bearing deletions were confirmed by genomic PCR. All strains were maintained on normal growth medium (NGM) plates at 20°C unless otherwise indicated, with E. coli strain OP50 as described (47).

**Table 1:** Worm strains used in the study.

| Name | Genotype |
| --- | --- |
| KX149 | ife-1(eu20[mKate::Myc3x::ife-1]) III, pgl-1(sam33[pgl-1::gfp::3xFlag]) IV |
| KX152 | ife-3(eu21[GFP::Flag3x::ife-3]) V |
| KX154 | ife-3(ok191)/eT1 III:V, him-8(e1489) IV |
| KX155 | ife-1(eu20[mKate2::Myc3x::ife-1]) III, ife-3(eu21[GFP::Flag3x::ife-3]) V |
| KX178 | ife-3(eu21[GFP::Flag3x::ife-3])/eT1 III:V, ife-1(bn127)/eT1 III:V, umnls9 [myo-2p::GFP] V |
| KX185 | ife-1(eu20[mKate2::Myc3x::ife-1]) III, ife-3(ok191)/eT1 III:V, him-8(e1489) IV |
| DUP64 | glh-1(sam24[glh-1::gfp::3xFlag]) I |
| DUP287 | glh-1(sam176[glh-1::wormScarlet(1-11)]) I |
| DG4222 | pos-1(tn1730[gfp::tev::3xflag::pos-1]) V |
| KX242 | ife-1(bn127) III, pos-1(tn1730[gfp::tev::3xflag::pos-1]) V |
| DG4739 | lin-41(tn1892[mScarlet::tev::3xflag::lin-41]) I, spn-4(tn1699[spn-4::gfp::3xflag]) V |
| KX243 | lin-41(tn1892[mScarlet::tev::3xflag::lin-41]) I, ife-1(bn127) III, spn-4(tn1699[spn-4::gfp::3xflag]) V |
| DG4269 | mex-3(tn1753[gfp::tev::3xflag::mex-3]) I |
| KX241 | mex-3(tn1753[gfp::tev::3xflag::mex-3]) I, ife-1(bn127) III |
| KX227 | glh-1(sam176[glh-1::wormScarlet(1-11)]) I, ife-3(eu21[GFP::Flag3x::ife-3]) V |
| LP306 | cpls53 [mex-5p::GFP-C1::PLC(delta)-PH::tbb-2 3'UTR + unc-119 (+)] II |
| LP307 | cpls54 [mex-5p::mKate::PLC(delta)-PH(A735T)::tbb-2 3'UTR + unc-119 (+)] II |
| CGC20 | dpy-18(e364)/eT1 III; unc-46(e177)/eT1, umnls9 [myo-2p::GFP + NeoR, III:9421936] V |
| N2 | +/+ (wild type) |
| KX223 | glh-1(sam24[glh-1::gfp::3xFlag]) I, ife-1(eu20[mKate2::Myc3x::ife-1]) III |
| RFK1257 | lotr-1(sam85[3xFLAG::gfp::lotr-1]) I, znfx-1(gg634[HA::tagRFP::znfx-1]) II |
| KX201 | znfx-1(gg634[HA::tagRFP::znfx-1]) II, ife-3(eu21[GFP::Flag3x::ife-3]) V |
| SS712 | ife-1(bn127) III |

**Table 2:**
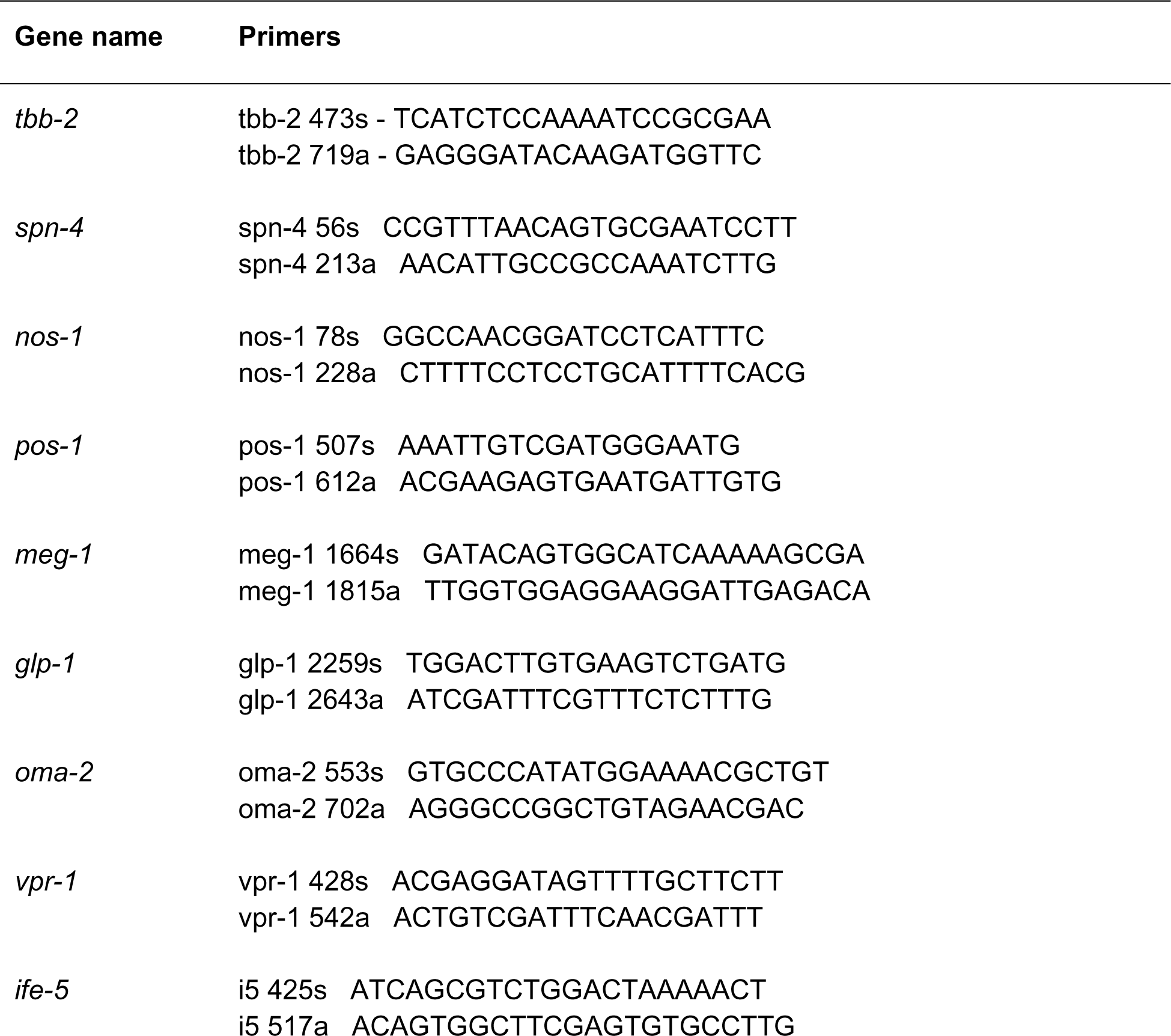
Primers used in this study.

### RNAi Treatment

L4 worms were fed *E. coli* strain HT115 harboring either control vector L4440 or gene-targeting construct (pT7-2::ife-1, ife-3, ifet-1, pgl-1/-3, or glh-1/-4) at 20C (except where noted) as described previously and F1 offspring were analyzed by microscopy following modest fixation and washing that preserves GFP, mKate2 and mScarlett fluorescence (40). To prepare RNAi feeding plates, frozen bacterial stock was streaked on 2x YT agar plates and grown overnight at 37°C. individual colonies were inoculated into 2x YT with carbenicillin 50ug/mL and grown for 16-19hrs on a shaker at 37°C. Cultures were pelleted by centrifugation and seeded on NGM plates coated with induction cocktail (1mM IPTG and carbenicillin 50 ug/mL) overnight at room temperature. L4 worms were placed onto RNAi plates at 20°C or 25°C and F1 adult off-spring were processed for imaging (confocal or standard fluorescence microscopy) as whole worms or dissected gonads.

### Microscopy and Immunostaining

For whole worms imaging, worms were mounted on 2% agar pads with levamisole 1mM and fluorescence images captured on a Zeiss Observer 7 or Axiovert 200M microscope on an HRM camera using Zen Blue software or Axiovision v4.5 (Carl Zeiss Imaging). For fluorescence quantification, image exposures (typically 0.5 to 10s), light intensity and camera sensitivity settings were fixed. Exposures in the Texas red and FITC green channels did not exceed 70% of the camera range. Fluorescence quantification was done using ImageJ 1.54F Fiji plugin and a macro created by F Bonnet at the MDIBL Light Microscopy Facility (see below). Colocalization quantification was done on ImageJ 1.54F colocalization plugin. Any image adjustments to contrast or brightness were conducted linearly. In all micrographs shown, mKate2 is pseudo-colored as magenta for individual fluorescence, but maintained as red in Merged images for red/green contrast (mKate vs GFP fusion proteins). GFP is always pseudo-colored green.

For native red, green fusion proteins, and DAPI fluorescence imaging, gonads were isolated from 1 day old gravid adults, gently fixed with 3% paraformaldehyde for 20 min on ice, washed twice in PTW followed by cold 70% ethanol as described in (40). Samples were imaged on a Zeiss Axiovert 200M or Observer 7 microscope on 2% agarose pads within 90 min after preparation. For gonad immunostaining, 3% PFA and 100% methanol-fixed gonads were stained with rabbit anti-IFET-1 (1:200) or anti-PGL-1 (1:1000) in PTWB overnight at 4°C (48). Gonads were washed three times at 10 min intervals in PTWB and then incubated in Alexa Fluor 568-conjugated goat anti-rabbit-IgG (1:400) secondary antibody (Invitrogen) in PTWB for 1 h at room temperature. The gonads were washed, stained with DAPI (0.05 ug/ml) and imaged as above using a 40x Neofluar air objective or 100x oil objective. Samples for confocal microscopy were prepared using identical gonad dissection and fixation. Confocal imaging was conducted using the Zeiss LSM 800 Microscope using a 63x water immersion objective. In experiments using fluorescence quantification, image exposures (typically 0.5 to 10s), light intensity and camera sensitivity settings were fixed. Exposures in the red and green fluorescence did not exceed 70% of the camera range. Fluorescence quantification was done using ImageJ 1.54F Fiji plugin and a macro created by F Bonnet at the MDIBL Light Microscopy Facility. Colocalization quantification was done on ImageJ 1.54F colocalization plugin. Any image adjustments to contrast or brightness were conducted linearly.

### High resolution microscopy

Young adult worms were mounted on 2% agar pads with levamisole (5mM) between slide and No.1.5 coverslip. Images were acquired using a point scanning confocal unit (LSM980, Carl Zeiss Microscopy, Germany) on a Zeiss Axio Examiner Z1 upright microscope stand (ref: 409000-9752-000) equipped with a Zeiss Plan-Apochromat 63x/1.4 Oil (ref:420782-9900-799) objective and Immersol 518F Immersion Oil (ref: 444960-0000-000, Carl Zeiss Microscopy, Germany). GFP and mKate/wrmScarlet fluorescence were excited with the 488 nm diode and with the 561 nm DPSS laser, respectively. Fluorescence was collected with Airyscan2 with the following detection wavelengths: GFP from 499 to 557 nm and mKate/wrmScarlet from 573-627 nm. Images were sequentially acquired in Super Resolution mode (SR) at zoom 10.6, with a line average of 1, a resolution of 244 x 274 pixels, 0.043 x 0.043 um/pixel, a pixel time of 0.64 us, in 16-bit, and in bidirectional mode. 2.0 um z-stack images were collected with a step size of 0.15 um with the Motorized Scanning Stage 130 x 85 PIEZO mounted on Z-PIEZO stage insert WSB500. The microscope was controlled using Zen Blue Software (Zen Pro 3.1), AiryScan images were processed using the “auto” mode and saved in CZI format. Fig 6H-K are 0.30 um maximum intensity projections of pachytene-stage germ cells in these living worms.

### Sample Preparation for LC-MS/MS

Worms were grown on 10 cm OP50 and chicken egg plates at 20°C then washed off plates using M9, allowed to purge gut contents for 30 min in M9. Worms were then floated over 35% sucrose at 4°C to remove debris and bacteria, washed with 0.1 M NaCL, then flash frozen in liquid nitrogen in the presence of 14 mM E64 protease inhibitor (Sigma-Aldrich), 4 mM Vanadyl-ribonucleoside RNase inhibitor (Sigma-Aldrich), and 0.2% Tween 20 detergent. Worm pellets were subsequently crushed in a mortar and pestle under liquid N2 with an equal volume of mRNP immunoprecipitation buffer (50 mM Tris-HCl [pH 8.0], 300 mM NaCl, 10 mM MgCl2, 1 mM EGTA, 2 mM dithiothreitol (DTT), 800 U of RNAsin/ml (Promega), 4 mM Vanadyl-ribonucleoside complex, 2X HALT complete protease inhibitor (ThermoFisher), 14 mM E64 protease inhibitor (SigmaAldrich), then centrifuged at 14,000 × g at 4°C for 15 min. The resultant high-speed supernatant was subjected to 20-50 μL of packed equilibrated GFP or RFP-targeted Nano-Trap agarose beads (Chromotek) and bound for 1.5 hrs. at 4°C while rotating. All immunoprecipitations were done on 6-10 mg of total C. elegans protein as measured by Coomassie spot staining relative to BSA standards. After binding, beads were washed 4X in high salt high detergent buffer (50 mM Tris-HCl [pH 7.8], 1 M NaCl, 0.5% NP-40, 0.5% deoxycholic acid), then additionally washed 3X in 20 mM Tris-HCl [pH 7.8]. The IP beads were sent for LC-MS/MS. Samples were subjected to on-bead tryptic digest, and desalted using C18 desalting spin columns (Pierce) per manufactures instructions. Each sample was analyzed in technical duplicate by LC-MS/MS using a Thermo Easy nLC 1200-QExactive HF. Collected data was then processed using Proteome Discoverer 2.1 by searching against the Uniprot *C. elegans* database, client specified database (IFE-1, IFE-3, tags), and a contaminants database, using Sequest. Results were further analyzed in Scaffold. Scaffold thresholds were set to: 2 peptide minimum, 95% protein and peptide scoring thresholds. Strains expressing the plextrin homology domain fused to the identical tags (mKate2::3xMyc::PH and GFP::3xFlag::PH) were also analyzed to control for proteins that bind either the fluorescent proteins or respective affinity tags. Fold Change (fc) ratios were calculated based on average spectral counts for both mKate2::IFE-1 and GFP::IFE-3 relative to their mKate2::PH and GFP::PH respective control precipitations. In cases where zero spectral (peptide) counts were identified in the control pulldown, a single spectral count (or 0.333 average) was arbitrarily assumed to allow divisibility. The assumption provided a minimal fold change that may be larger. Proteins with an average of 4 spectral count in two technical replicates and fold changes of >2 with a p-value <0.1 were considered potential interactors. These conditions were used to generate the distinct lists described for IFE-1 and IFE-3.

### RNA-Seq Analysis on Immunoprecipitates (RIP-Seq)

Samples were prepared identically to those for LC-MS/MS except that the wash buffer contained 0.3M NaCl and no secondary wash was performed. (We observed that yields of RNA were severely depleted using 1M NaCl to wash.) The Trap beads (or an aliquot of the initial homogenate for unenriched control) were extracted with Trizol (Life Technologies) according to the manufacturer’s protocol. Total RNA was also isolated from the same frozen worm samples by standard methods using Trizol (40). Supernatant recovery was followed by 0.7 volume isopropanol precipitation. Resuspended precipitate was further extracted with phenol:chloroform:isoamyl alcohol (25:24:1), and twice with chloroform:isoamyl alcohol (24:1), then ethanol precipitated. GlycoBlue (Invitrogen) was used as a co-precipitant. RNA quality and quantity was determined using NanoDrop® ND-1000 spectrophotometer. RNA sequencing services were outsourced to Admera Health, LLC.

Sequence reads of each sample were pseudo-aligned to *C. elegans* Ensemble reference transcriptome (WBcel235) and the gene transcript abundance was quantified by using Kallisto (v.0.48.0). Differential expression of genes and transcripts were achieved by using DESeq2 (v.1.34.0) in RStudio (Build 386 with R v.4.4.1) with a significance set at adjusted p-value<0.05 (green & purple in Volcano plot) and log2FoldChange (fc) larger than 1 (purple). Functional analyses were conducted using ClusterProfiler (v4.2.2) and a gene-set enrichment assay. Note that IFE-3 mRNPs retained more mRNAs than IFE-1 mRNPs, even wh.en normalized for mKate2::3xMyc or GFP::3xFlag controls (data not shown)

### Western blotting of proteins from IP for validation

IFE-1 and IFE-3 Trap IP beads were boiled in 4X SDS load buffer for 3 min, and 30 μg of protein for each sample was resolved on 10% SDS PAGE gels. Protein content was determined by spot assay and Coomassie Brilliant Blue staining vs BSA standards. Protein from the gel was transferred to polyvinylidene fluoride (PVDF) membranes overnight (70 mA) and membrate blocked with TST (10 mM Tris-HCl ph 7.4, 150 mM NaCl, 0.05% Tween 20) containing 5% dry non-fat milk and 1% BSA for 1 hr. at room temperature. Membrane was then incubated with either a 1:4000 dilution of a GFP specific primary antibody (Sigma; G1544) or a 1:3000 dilution of a RFP specific primary antibody (Invitrogen; R10367) in TST containing 5% dry non-fat milk and 1% BSA for 1 hr. at room temperature. Blots were washed 5 times in TST and incubated with a 1:15,000 dilution of goat anti-rabbit IgG secondary antibody conjugated to horseradish peroxidase (HRP) in TST containing 5% dry non-fat milk and 1% BSA for 1 hr. at room temperature, then washed extensively. Detection was done by ECL+ kit (ThermoFisher) per manufactures instructions. Image acquisition was done using a Amersham Typhoon Fluorescence/Phosphorimaging scanner (GE Healthcare) at the ECU PhIFI Core Facility.

### qRT-PCR on IP samples

For qPCR analysis of IFE-1 and IFE-3 IP samples and total RNA was used. cDNA from both RNA-IP sample, 20ul was synthesized using iScript Advanced cDNA synthesis kit (Bio-Rad Laboratories) according to the manufacturer’s instructions. qRT-PCR was performed in triplicate on a OPUS CFX-96 Real-Time System (Bio-Rad Laboratories) using Sso Fast Evagreen Super mix (Bio-Rad Laboratories), according to the manufacturer’s instructions. Annealing temperatures and input cDNA amounts were determined using standard efficiency titrations (40,44). Primer sequences include: tbb-2 (forward 5’- TCATCTCCAAAATCCGCGAA -3’, reverse 5’- GAGGGATACAAGATGGTTC -3’); spn-4 (forward 5’- CCGTTTAACAGTGCGAATCCTT -3’, reverse 5’- AACATTGCCGCCAAATCTTG -3’); nos-1 (forward 5’- GGCCAACGGATCCTCATTTC -3’, reverse 5’- CTTTTCCTCCTGCATTTTCACG -3’); pos-1 (forward 5’- AAATTGTCGATGGGAATG -3’, reverse 5’- ACGAAGAGTGAATGATTGTG -3’); meg-1 (forward 5’- GATACAGTGGCATCAAAAAGCGA -3’, reverse 5’- TTGGTGGAGGAAGGATTGAGACA -3’); glp-1(forward 5’- TGGACTTGTGAAGTCTGATG -3’, reverse 5’- ATCGATTTCGTTTCTCTTTG -3’); oma-2 (forward 5’- GTGCCCATATGGAAAACGCTGT -3’, reverse 5’- AGGGCCGGCTGTAGAACGAC -3’); and vpr-1 (forward 5’- ACGAGGATAGTTTTGCTTCTT -3’, reverse 5’- ACTGTCGATTTCAACGATTT -3’).

## Results

### IFE-1 and IFE-3 localization in germ cell granules is re-established in the embryo

The functions of eIF4E isoforms in the germline of *C. elegans* are simply the canonical roles of mRNA cap-binding proteins. Rather, our work and others have shown that they can act both positively and negatively for mRNA translation initiation (12,24), and their activities are mRNA selective (40–44). We showed that the two predominant germline-enriched eIF4Es, IFE-1 and IFE-3, are differentially expressed in the late larval to adult hermaphrodite gonad during the transition from spermatogenesis to oogenesis (40). IFE-1 predominates in early, undifferentiated germ cells and sperm, while IFE-3 accumulates to become the predominant isoform in oocytes. The distinct expression and localization patterns are obvious in dual red/green fluorescent worms that were CRISPR/Cas9 edited at the *ife-1* and *ife-3* gene loci to create in-frame fusions: IFE-1::mKate2 and IFE-3::GFP (Fig. 1). Both IFE-1 and IFE-3 are present in the distal germ cells, from the mitotic zone through diakinesis both as soluble cytoplasmic factors and in germ granule condensates (Fig. 1A and Huggins, 2020 #1198}. As germ granules dissipate in late spermatocytes and during oocyte maturation, both eIF4Es become fully soluble in their respective gametes. Here we show that IFE-1 and IFE-3 are again abundant in early embryos, re-establishing positions as adjacent but non-overlapping puncta in blastomeres of the P lineage (Fig. 1A, insets). Association of IFE-1 and IFE-3 with germ granules is due to interactions with their 4EIPs, PGL-1 and IFET-1, respectively (40,46,49). The separate localization is indicative that each eIF4E assembles into independent mRNPs not only in germ cells and gametes but also in embryos.

**Figure 1.**
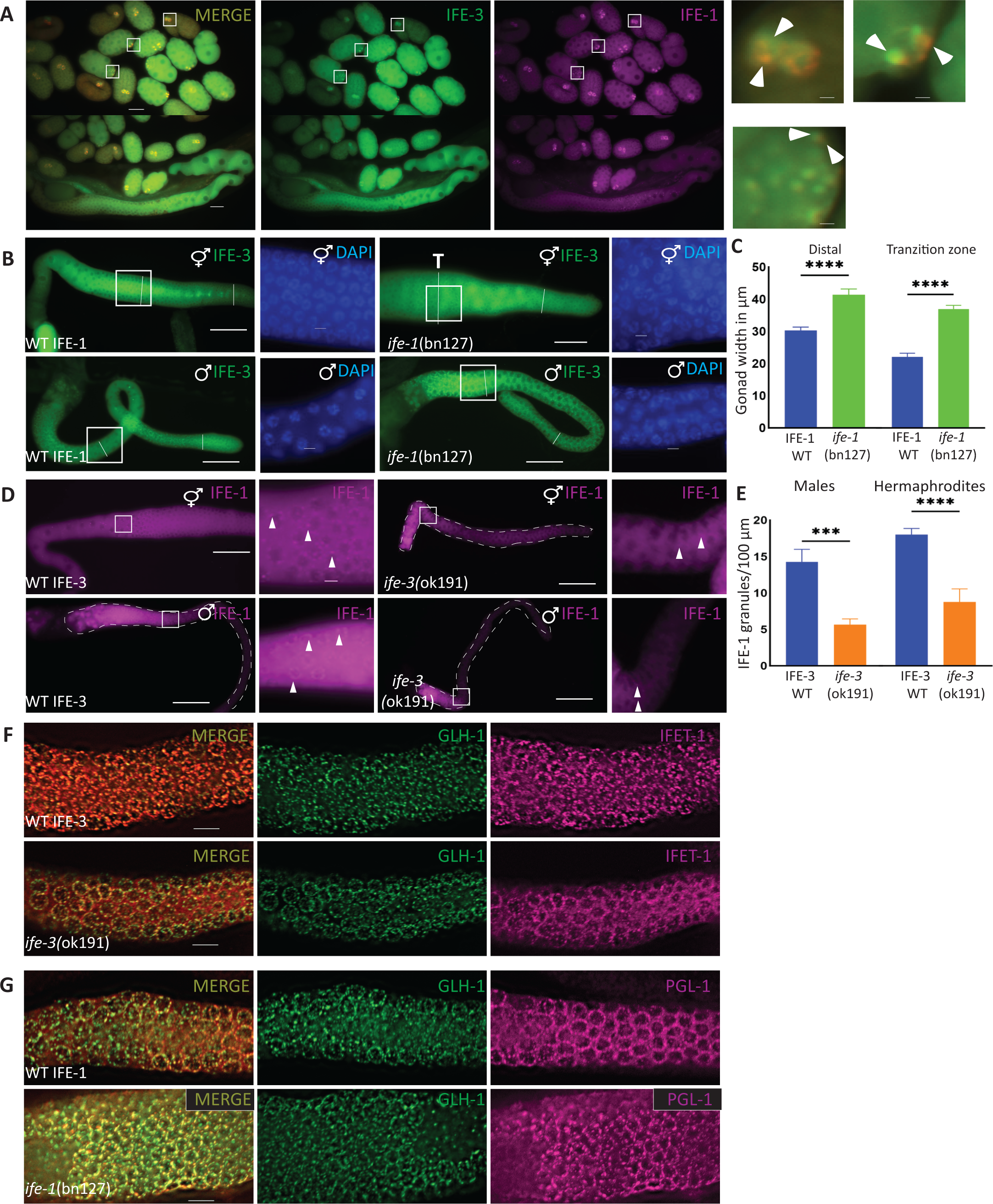
IFE-1 and IFE-3 localize differently in germ cell granules but have little effect on cognate 4EIPs. IFE-1 and IFE-3 are adjacent but not fully overlapping in early embryo P granules. A. IFE-1 and IFE-3 localization pattern in embryos in the CRISPR/Cas9 dual fluorescent mKate2:: 3xMyc::IFE-1 and GFP:: 3xFlag::IFE-3 strain. Insets (white boxes) show 10-fold magnification of germ granules of comma stage, ∼30 cell and 8 cell stages, respectively. Loss of IFE-1 or IFE-3 change phenotype, but neither affects the localization of other germ granule proteins. B. Gonads of IFE-1 null worms [*ife-1*(bn127)] showing GFP::IFE-3 expression. Hermaphrodite gonads were enlarged, and nuclear organization disrupted into the rachis; no similar change was seen in male gonads. C. Gonad width quantification (hermaphrodites only) at both distal and transition zone (T). White lines show measurement positions. D. IFE-3 null [*ife-3*(ok191)] causes a masculinization phenotype in hermaphrodites and reduced mKate::IFE-1 in granules. E. IFE-1 granules quantified per square 100µm in both males and hermaphrodites. F,G. Localization of other P granule proteins was unperturbed in either *ife-1*(bn127) null or *ife-3*(ok191) null gonads. F. IFET-1 immunostaining in GLH-1::GFP background in wildtype and *ife-3*(ok191) worm gonads. G. PGL-1 immunostaining in GLH-1::GFP background in wildtype versus *ife-1*(bn127) gonads. mKate2 is pseudo-colored as magenta for individual fluorescence but maintained as red in Merged images for red/green contrast. D – Distal gonad; T – Transition zone gonad. Scale bar: 20µm for A-D, 5µm for F, G and insets. For C and E, n>10; p values: **** >0.0001, *** >0.001, SEM for error bars, p value 0.05.

The expression patterns of IFE-1 and IFE-3 vary inversely later in germline development, particularly as meiotic cells differentiate into sperm or oocytes, in a way that coincides with the respective phenotypes of IFE-3 and IFE-1 null mutations (40,46). Although both show moderate expression in the germline of both hermaphrodites and males, IFE-1 levels peak in primary and secondary spermatocytes but remains low in oocytes (Fig. 1A-D). In the oogenic gonad, IFE-3 increases from the transition zone through proximal region and accumulates in growing oocytes through the −1 position (Fig. 1A, lower panels). During this period IFE-1 dissipates (but is not lost) in the hermaphrodite proximal gonad. Much like other pro-spermatogenic proteins (except MSPs), IFE-1 is relegated to the residual bodies (RBs) upon spermatid budding (Fig. 1D). Curiously following fertilization, both proteins are again abundant, primarily in soluble form except in the P lineage cells, where condensation in germ granules can again be observed (Fig 1A, white boxes). However, the red/green dual fluorescence indicates that IFE-3 decreases in late blastulae (>50 cell), and IFE-1 is again the predominant eIF4E already in comma-staged embryos (Fig 1A, first panel).

### Proper localization of each germline IFE isoform is dependent on the other isoform

Our work has previously shown that IFE-1 loss arrests late spermatocyte maturation and mis-regulates oocyte maturation (41,44), whereas loss of IFE-3 results in impaired germline sex determination (sperm to oocyte switch) and causes embryonic lethality (40). Biochemical analysis suggested that the two IFEs may be co-isolated (see Fig. 3), so we now asked was if the presence of one eIF4E isoform affected the correct localization of the other isoform. We addressed both the developmental germline abnormalities and any changes in their localization patterns using worms bearing deletions in either the *ife-1* or *ife-3* gene and the fluorescent-tagged gene for the other isoform. Changes in gonad morphology were observed in dissected male and hermaphrodite gonads that lacked IFE-1 or IFE-3. Gonad width was significantly increased in the distal and the transition zone gonad of *ife-1*(bn127) mutants (Fig. 1B, C, n=7), but IFE-3 expression was unperturbed. Loss of IFE-1 caused nuclei to be displaced to the rachis coincident with gonad enlargement. IFE-3 localization to germ granules was not affected by IFE-1 loss but was curiously disrupted along the rachis lattice-like structures (40) compared to wildtype worms. Under these conditions distal germ nuclei apparently fall from their peripheral arrangement along the sheath and mis-localize into the rachis. A similar nuclear displacement was observed by others in the absence of IFET-1 [(48) and Fig. 2F], but its loss also displaces IFE-3 from germ granules (40). Thus both IFE-1 and IFET-1 play roles in gonad architecture.

**Figure 2.**
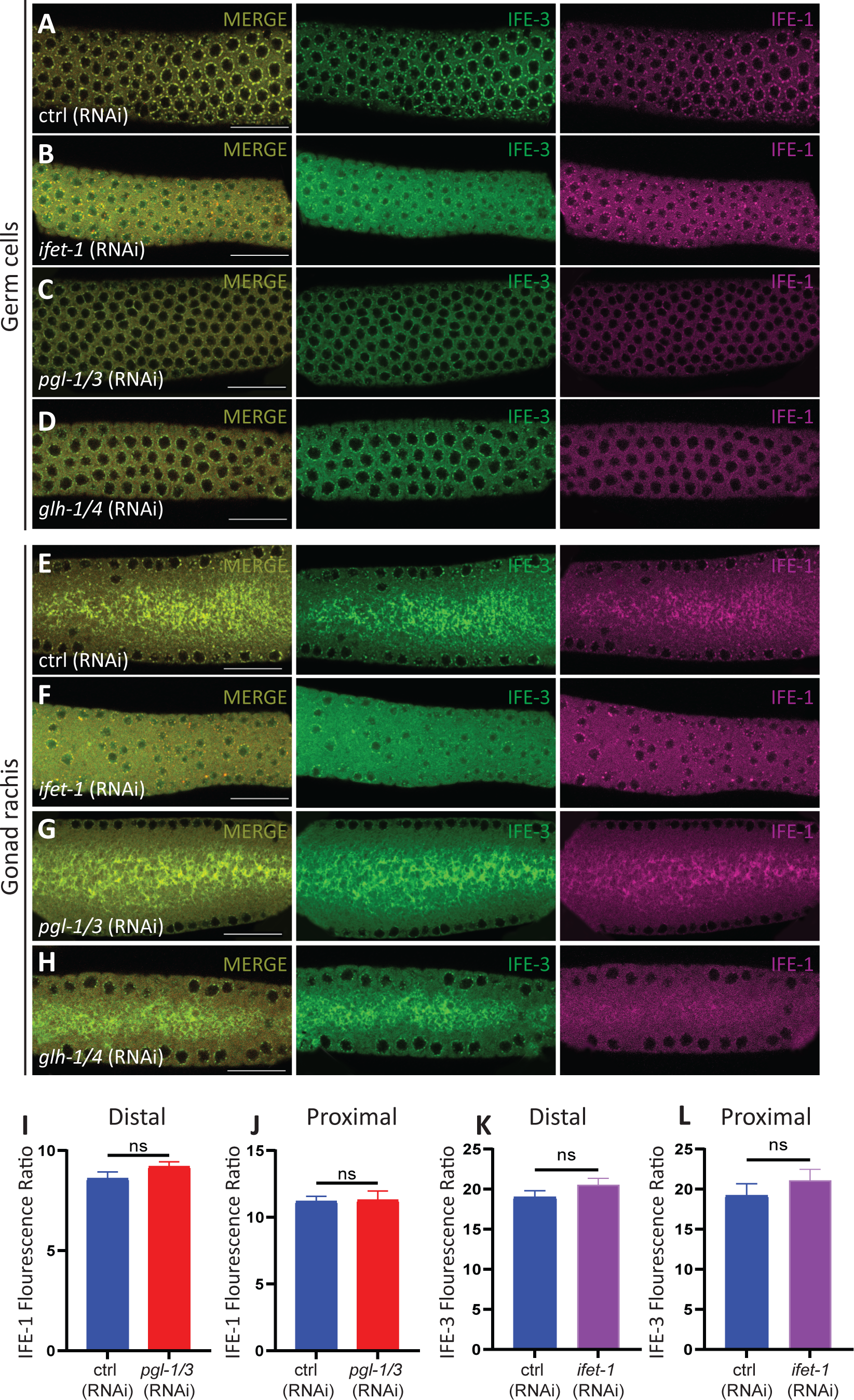
Loss of 4EIPs or GLH-1 effect nuclear arrangement and IFE localizations. Confocal microscopy on gonads dissected from worms bearing both CRISPR/Cas9 fluorescently-tagged mKate2::TEV::3xmyc::IFE-1 and GFP::TEV::3xflag::IFE-3 are shown. Focal planes at the top of the distal region of gonad were used to evaluate distribution IFE-1 and IFE-3 in germ cells (A-D, I, K) and cross-sectional planes in more distal regions of the same gonads used to evaluate the rachis morphology (E-H, J, L). Worms had been treated by fed RNAi for A, E, ctrl(RNAi); B, F, *ifet-1*(RNAi); C, G, *pgl-1/3*(RNAi); D, H, *glh-1/4*(RNAi). Graphs show remaining total mKate::IFE-1 fluorescence (I and J) or GFP::IFE-3 fluorescence in the distal and proximal dissected gonads (n>15) after RNAi treatment for control (ctrl), *pgl-1*, or *ifet-1* (K and L). ns – not significant using unpaired t test, p value >0.05, error bars show SEM; scale bar: 20µm.

**Figure 3:**
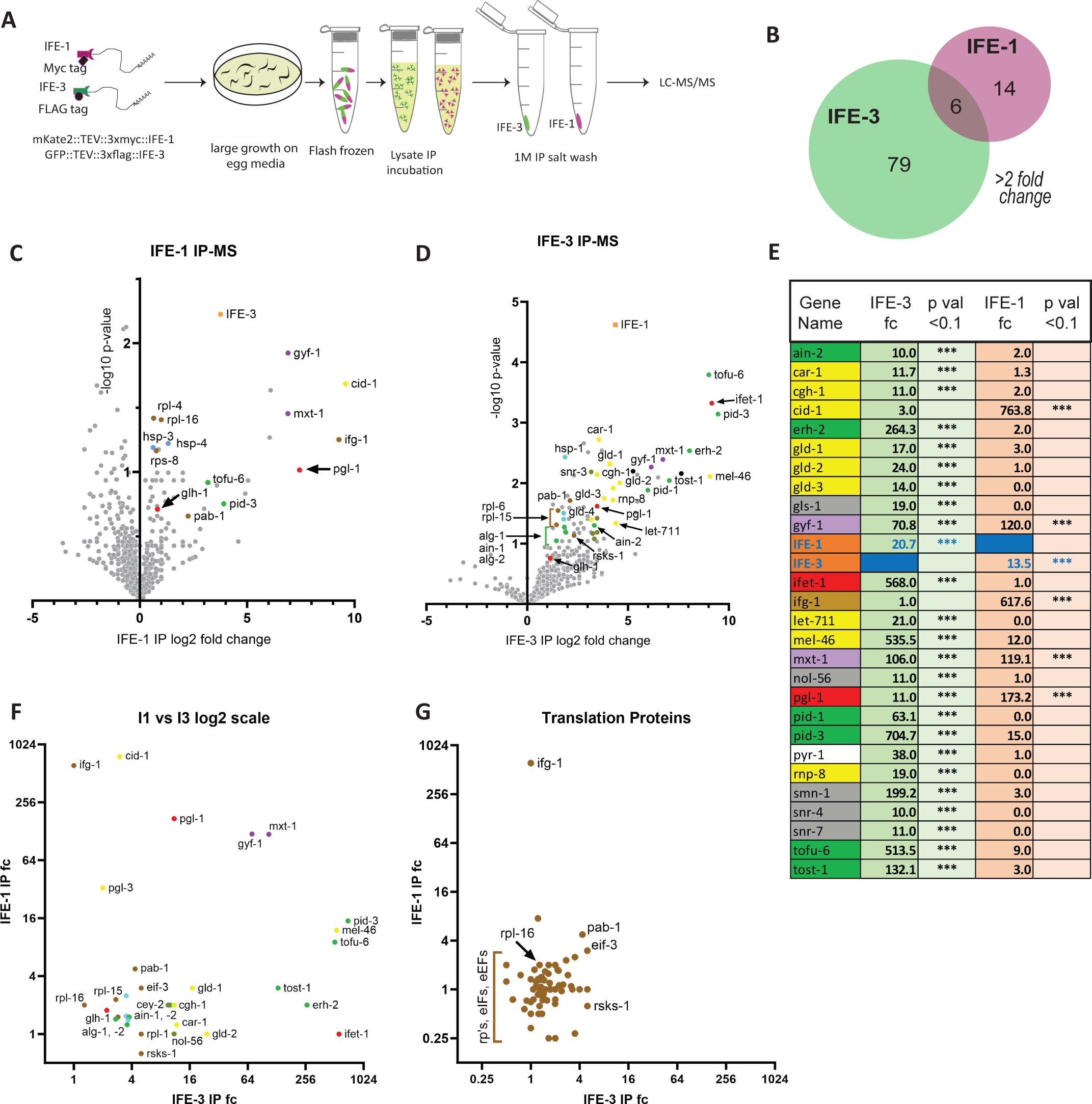
Proteomic identification of proteins in the IFE-1 and IFE-3 mRNPs. A. Schematic shows steps in sample preparation for LC-MS; details are found in Methods. B. A Venn diagram of the number of proteins enriched at least two-fold with either IFE-1 or IFE-3 shows that of the 99 identified proteins, only 6 were found in both mRNPs. C, D. Volcano plots showing log2-fold enrichment versus the -log10 of the p value for the IFE-1 (C) and IFE-3 (D) mRNPs compared to the initial total protein lysate. Gray dots indicate all proteins identified by at least 4 peptides. Color coding of dots shows proteins of known function and is consistent in C-G. Binding of each IFE in the other’s mRNP is shown in orange. 4EIPs bound directly to IFE-1 or −3 are in red dots, as is the Vasa helicase, GLH-1. Indiscriminate 4EIPs are shown by purple dots. Dots also show relative binding of germline RBPs, poly(A) polymerases and helicases (yellow), Argonaut-associated and PETISCO small RNA-related proteins (green), small nuclear RNA-pathway proteins (dark gray), translation-related proteins (brown). E. Complete list of the 28 most tightly bound proteins (*** indicates >10-fold enriched, p <0.1) from named genes in WormBase show only one unrelated protein (white box). 26 are highly enriched with IFE-3. Only 6 are highly enriched with IFE-1, of which only 2 are exclusive; IFG-1 and CID-1. PGL-1 is 16-fold better enriched with IFE-1. F. Direct comparison (log2 scale) of moderate and highly enriched proteins with known functions for IFE-1 (I1) and IFE-3 (I3). G. Since IFE-1 and IFE-3 are translation initiation factors, a direct comparison (log2 scale) of the relative binding of identified ribosomal proteins (rp’s) and other translation initiation (eIFs) or elongation factors (eEFs). Most are just 2-8-fold enriched by either eIF4E. Only proteins with greater enrichment and p<0.1 are labeled. fc: fold change.

The primary developmental role of IFE-3 is to facilitate the sperm to oocyte switch and sustain oocyte fate (40). Masculinization of the hermaphrodite gonad caused by the *ife-3*(ok191) mutation was apparent by enhanced IFE-1 levels in resulting spermatocytes (Fig. 1D). There are very few developing oocytes and most nuclei are entering a spermatogenic fate. Interestingly, the IFE-1 localization to germ granules is diminished in the *ife-3* deletion strain by 57% in males and 55% in hermaphrodites (Fig. 1E, n=7), even though IFE-1 becomes induced normally in male germ cells (Fig. 1D). These results show that each eIF4E isoform has a unique role in germ cell fate and gonad organization. However, their roles are not fully independent. IFE-1 plays some role in IFE-3 association with the rachis lattice and IFE-3 has a role in maintaining IFE-1 in the germ granule.

We posed the same question for the localization of each IFE’s cognate binding partner: Do the 4EIPs or other integral germ granule components rely on IFE-1 or IFE-3 for localization, particularly when germline development is impacted? Using GLH-1::GFP as a central marker for germ granule structure, we assessed IFET-1 localization (the IFE-3-binding partner) in the absence of IFE-3, and PGL-1 localization (the IFE-1 binding partner) in the absence of IFE-1, by immunostaining for each 4EIP (Fig. 1F, G). Both 4EIPs remained strongly associated with germ granules without their cognate eIF4Es; IFET-1 localized with GLH-1 even without IFE-3 (Fig. 1F) and PGL-1 localized with GLH-1 even without IFE-1 (Fig. 1G). Thus, the presence of one IFE may affect the localization of the other isoform in certain structures. Previous studies showed that PGL-1 and IFET-1 are conversely essential to localizing IFE-1 and IFE-3, respectively (40). But neither IFE-1 nor IFE-3 is required for localization of germ granule components more integral to the condensate. It is worth noting that IFE-5 protein is 80% identical to IFE-1, but it does not bind PGL-1 nor does it functionally substitute for *ife-1* loss (29). As a precaution we determined that *ife-5* gene expression is not upregulated in various mutant strains (Suppl Fig. 4). These findings establish an observable hierarchy in the protein binding layers within germ granule substructures. The eIF4E isoforms are clearly the most “external” in this hierarchy, yet still influence one another. The following experiments seek to determine the links in the IFE-1/IFE-3 relationship.

### 4EIPs affect the localization but not levels of IFE-1 and IFE-3

The *C. elegans* germline eIF4Es are sub-localized within germ granule structures by their interacting partners, 4EIPs (12,40). The IFE-1:PGL-1 and IFE-3:IFET-1 complexes have been shown to localize distinctly in the germ granule (40,46). There is also significant IFE-3 in lattice structures in the rachis that co-localizes with IFET-1 (49Sengupta, 2013 #960). Numerous studies have shown that loss of PGL-1, GLH-1, or IFET-1 disrupts germ cell development leading to infertility (22,40,48,50–52). With a better understanding of the hierarchy of germ granule protein interactions described above, we can now determine how the overall expression or distribution IFE-1 and IFE-3 are impacted by removal of the 4EIPs and the Vasa homolog, GLH-1. Particularly in the distal gonad, the overall IFE-1 and IFE-3 expression levels are not changed by the loss of IFET-1, PGL-1 or GLH-1 (Fig. 2). However, their respective localization to the P granules (as well as P granule size) is impacted in very particular ways. Confocal imaging of GFP::IFE-3 and mKate2::IFE-1 upon *ifet-1*(RNAi) showed increase in the germ granule size for both IFE-1 and IFE-3 labeling and a non-regular, mildly clumped distribution around the nuclear periphery (Fig 2B). Much of the IFE-3 association was lost from germ granules, as it was from the rachis lattice (Fig. 2F). Indeed, the lattice-like structure in the syncytial cytoplasm disappeared and germ cell nuclei reside aberrantly in the rachis without IFET-1. The loss of lattice-like structure also allowed a more even distribution of the soluble IFE-3 throughout the rachis. More granules appeared to have only IFE-1 present, suggesting its proportion in germ granule versus soluble was not changed (Fig. 2B, F). Similar *pgl-1*(RNAi)-treated germlines routinely showed a tighter arrangement of germ cell nuclei in the gonad. Somewhat unexpectedly, both IFE-1 and IFE-3 are lost from P granules (Fig. 2C, G). No release of IFE-3 from rachis lattice was observed upon PGL-1 depletion. Finally, depletion of the Vasa homologs, GLH-1 and GLH-4) caused a substantial loss of IFE-1, and to some extent IFE-3, from germ granules (Fig. 2D, H). This is consistent with the direct interaction of PGL-1 with GLH-1, and thus supports the hierarchical binding arrangement of GLH-1:PGL-1:IFE-1 and GLH-1::PGL-1::IFET-1::IFE-3. Careful quantification of fluorescence intensity indicated there was no significant loss of IFE-1 nor IFE-3 abundance after these RNAi depletions (Fig. 2I-L), nor any change in the IFE-1 to IFE-3 ratio. The results indicate that some remodeling of proteins in the germ cell occurs occurs as availability of these interacting components is altered.

### IFE-1 and IFE-3 form different mRNP complexes with overlapping secondary interactions

It is intriguing that a portion of both major eIF4Es are localized germ granules, but are adjacent rather than overlapping and held in place by unrelated 4EIPs (40). More puzzling is the fact that loss of IFE-1 disrupts translation for late sperm/oocyte development, while loss of IFE-3 affects primarily germline sex-determining translational control and much later embryonic development. To gain insight into how these mRNA cap-binding isoforms may play roles in such divergent development processes, we undertook both proteomic and mRNA analysis of their respective mRNP complexes. Since eIF4Es have been found in both active translation initiation complexes and repressed, translationally inert complexes (5,12), we expected to find evidence of both activities. The IFEs share a high degree of homology (29), so we took advantage of the 3xMyc and 3xFlag tags on the fluorescently labeled IFE-1 and IFE-3 to affinity purify each isoform from a single worm strain expressing both tagged isoforms. Immunoprecipitations were subjected to high stringency salt wash, but not nuclease, to isolate complete mRNP structures. Retained proteins were compared to control precipitations from strains expressing the fluorescence and affinity tags but lacking IFE sequences (see Methods). This allowed identification of interacting proteins and mRNAs by LC-MS/MS (IP-MS) and RNA-Seq (RIP-Seq), respectively in intact complexes (Fig. 3A). We identified 20 proteins associated with IFE-1 (>2-fold enriched, p<0.1) and 85 proteins associated with IFE-3 under those same conditions (Fig. 3B). Volcano plots show a larger subset of binding partners for IFE-3 relative to its GFP::3xFlag::PH control than for IFE-1 relative to its mKate2::3xMyc::PH control (Fig. 3C,D). We cannot rule out that our capture of IFE-1 was less efficient than IFE-3, however western blotting indicated similar yields of each tagged IFE (Suppl Fig. 2). Perhaps not surprisingly, the majority of captured proteins were of known germline function, either as 4EIPs and their direct partners (Fig. 3 C-E, red and purple points), members of the PETISCO or Argonaut small RNA complexes (green), translation factors/ribosomal proteins (brown), or germline helicases, RNA binding proteins, translational repressors and P body components (yellow). Perhaps most striking was that IFE-1 and IFE-3 bound one another under the conditions of mRNP isolation (more than 10-fold enrichment). This gives further credence to linked but separable roles for each IFE suggested by their non-overlapping but adjacent localization within P granule substructures (Fig. 1A, Fig. 2 and (40). Yet it is also clear from the IP-MS data that the constellation of IFE-1- and IFE-3-interactors are quite different.

By focusing more stringently on proteins at least ten-fold enriched we hoped to gain an understanding of functional differences between IFE-1 and IFE-3. The list (Fig. 3E) identified nearly exclusively proteins that have known roles in RNA regulation in the germline. It became clear that each eIF4E isoform preferentially binds a more defined set of partners that are non-overlapping. Only three proteins are common to both. MXT-1 (ortholog of vertebrate Mextli) and GYF-1 (ortholog of human Parkinson’s protein GIGYF2) are known 4E-IPs that bind to multiple somatic eIF4E isoforms (53,54). These were roughly equally bound by IFE-1 and IFE-3, providing further evidence that they are both universal 4E-IPs. Interestingly, the protein KH-domain germ granule protein PGL-1 was identified in both pull-down sets, though it bound more efficiently to IFE-1 (173-fold enrichment) than to IFE-3 (11-fold enrichment). PGL-1 is the cognate 4E-IP for IFE-1 and our early studies demonstrated its direct interaction with IFE-1 but not with IFE-3 or other eIF4E isoforms (46). However, IFE-3 likely interacts indirectly with PGL-1 through its cognate 4E-IP, IFET-1, as shown by others (48,49). The table in Fig. 3E directly compares the strength of binding (by fold enrichment) of major proteins identified by LC-MS, and spatially depicts the preferential mRNP compositions for each IFE. One of the two proteins most enriched by IFE-1 is IFG-1, the eIF4G ortholog in *C. elegans*. The long isoform (p170) of IFG-1 promotes mRNA cap-dependent translation initiation events by association with eIF4Es (55,56).CID-1 (also called CDE-1 or PUP-1) encodes a poly(U)polymerase and is strongly bound by IFE-1 (57–60). The relationship of 3’ poly(U) elongation of RNAs to their 5’ cap-binding is curiously unknown. By contrast to IFE-1, the proteins most strongly associated with IFE-3 are proteins with largely RNA repressor functions. These include its cognate 4E-IP IFET-1, shown by others to bind IFE-3 directly (49), the germline helicase MEL-46 (DDX20 helicase homologue), and all of the factors identified in the PETISCO complex (PID-1, PID-3, TOFU-6, ERH-2, TOST-1). PETISCO is an essential embryonic biosynthetic complex involved in 21U small RNA processing that was previously shown to require IFE-3 (61,62). Other RNA-binding proteins that act in translational repression (GLD-1, RNP-8), or P body/degradative functions (CAR-1, CGH-1, LET-711) were also substantially enriched in IFE-3 mRNPs, strongly suggesting a role in mRNA repression for this isoform. The generally repressive nature of IFE-3 interactors, in contrast to the positive translational interaction of IFE-1 and IFG-1, suggests that these eIF4E isoforms are not equally active in germ cells to recruitment mRNAs to ribosomes. However, initiation complexes and ribosome associations are transient and dynamic such that they rarely accumulate stably in the absence of inhibitors (4,5,7,8,43,63). We addressed the relative abundance of all translation and ribosome-related proteins captured with IFE-1 and IFE-3 (Fig. 3G). Most were enriched only roughly twofold, and just a few ribosomal proteins reached statistical significance (p<0.1). INF-1 (eIF4A, also part of eIF4 complex) was not enriched by either eIF4E, again consistent with transient interaction. However, IFG-1 (eIF4G) was enriched more than 600-fold with IFE-1, suggesting strong association in the productive initiation complex. PAB-1 was enriched 4-fold with both IFE-1 and IFE-3, but did not reach significance (p>0.2). Instead, one subunit of eIF3 (eif-3.G) and the rpS6 kinase RSKS-1 were each 5-fold enriched with IFE-3. Translation initiation factor eIF3 has been shown to bind the Vasa homolog GLH-1 that is clearly integral to mRNA regulation by germ granules (22). The germline polyadenylation proteins GLD-2 and −3 proteins were 14-24-fold enriched in the IFE-3 mRNP. These proteins all have known roles in germline stem cells (see Discussion).

One unexpected set of proteins associated with IFE-3 that have suspected nuclear/nucleolar functions. Most significantly, SMN-1 (survival of motor neurons-1) was nearly 200-fold enriched in IFE-3 mRNPs. SMN-1 participates in snRNP (snRNP small nuclear ribonuclear proteins) homeostasis to regulate transcripts at the level of pre-mRNA splicing (64). Other binding partners with nuclear functions include germline primary mRNA 3’-end processing protein GLS-1, nuclear protein 56 (NOL-56) and small nuclear RNPs Sm and G (SNR-4, −7, Fig. 3E, gray shaded). Overall, the uniquely different mRNP composition for each IFE suggest they participate in different functional RNA processes during germ cell development and embryogenesis. Given that both exist in separable substructures of the germ granule, as well as free in the cytoplasm, each IFE may have multiple roles (that presuppose multiple mRNP types) in mRNA trafficking, translation, and turnover.

### IFE-1 and IFE-3 mRNPs enrich different mRNA populations

To identify mRNAs that reside in the IFE mRNPs, we characterized the mRNA population captured by a similar immunoprecipitation strategy as described above (RIP-Seq, Fig. 4A). We postulated that “cargo” mRNAs in each of the IFE mRNPs may have some relationship to the mRNAs populations we have already shown to be translationally regulated by IFE-1 (41,44) and IFE-3 (40). By initially comparing the mRNA populations enriched by pulldown vs the control population (total RNA), we identified how many identified mRNAs were significantly enriched from total RNA in the IFE-1 pulldown and the IFE-3 pulldown. It should be noted that RNA-Seq reads are compiled from equal amounts of precipitated RNA and the total RNA sample (input) derived from the dual IFE-tagged strain. Therefore, differences reflect actual enrichment relative to the general mRNA population (Fig. 4B, C). Using a minimum enrichment of two-fold and p-value <0.05, there were 493 mRNAs enriched in the IFE-1 mRNP, whereas 409 mRNAs were enriched in the IFE-3 mRNP (volcano plot, purple dots right of Y axis Fig. 4B,C) using the same parameters. Most purple dots (except those described below) were unassigned gene names and unknown function in WormBase. Exceptions of very highly enriched, identified sequences were abundant housekeeping mRNAs transcripts, almost exclusively ribosomal protein (rp) mRNAs for IFE-1 and histone (his) mRNAs for IFE-3 (Supplemental Fig. 1G, H). More intriguing was direct comparison of the IFE-1 mRNP pulldown to the IFE-3 mRNP pulldown. The disparity of mRNA cargo identified by RIP-Seq was clearly evident, showing that IFE-1 clearly does not retain as much mRNA as IFE-3 (Fig 4D, both green and purple points had adjusted p value of <0.05). Direct comparison identified 219 mRNAs were at least two-fold more abundant in the IFE-3 mRNP than in the IFE-1 mRNP (volcano plot, Fig4D, purple), only 101 bound IFE-1 better. When we considered just annotated genes, IFE-3 was found to bind 109 identified mRNAs including a substantial number of translationally controlled critical germline mRNAs (see below) and 40 histone mRNAs (data not shown). By contrast, the 39 IFE-1-retained mRNAs encoded metabolic enzymes and structural proteins (data not shown), none of which had vital germline functions. Not surprisingly, a gene ratio analysis of mRNAs enriched by IFE-1 or IFE-3 showed products similarly involved in “peptide metabolic” and “amide biosynthetic” processes (protein synthesis), cell cycle, reproduction, germ cell development, gamete generation, oogenesis, and mitotic processes (Suppl Fig. 1). The assignments are consistent with regulatory roles in translation for IFE-1 and IFE-3 in early developmental metabolism and proliferation. More interestingly, the gene ratio analysis for mRNAs binding preferentially to IFE-3 over IFE-1 indicated a focus on reproduction, cell cycle, oogenesis, negative regulation of translation and embryonic pattern specification (Suppl Fig. 1C,F). These are all biological functions are substantiated by our previous phenotypic and translational analyses of *ife-3* (29,40).

**Figure 4:**
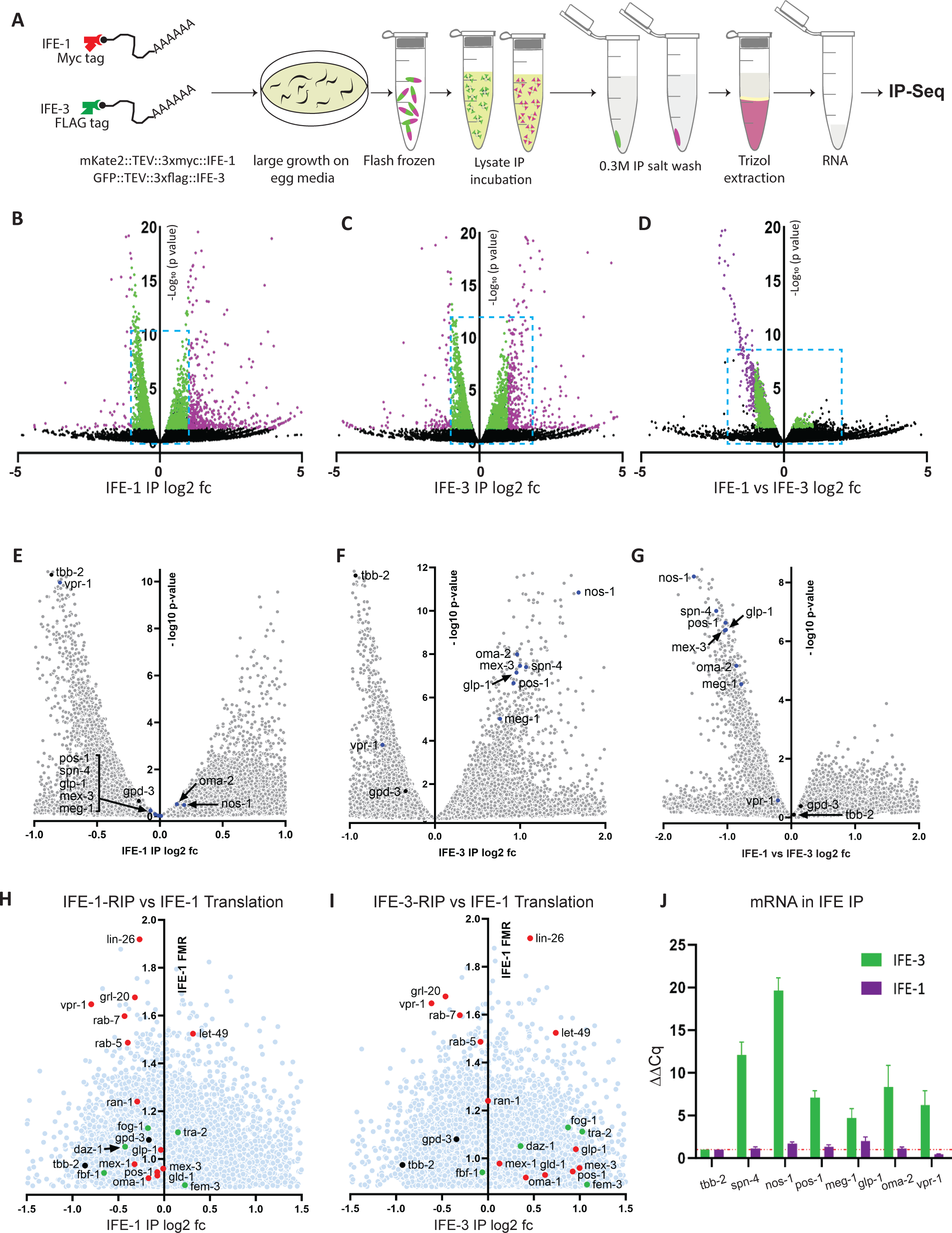
RIP-Seq identification of mRNAs in the IFE-1 and IFE-3 mRNPs. A. Schematic shows steps in sample preparation for RIP-Seq; details are found in Methods. Volcano plots comparing RNA Seq findings from (B) IFE-1 IP versus total lysate RNA, (C) IFE-3 IP versus total RNA, and (D) direct comparison of the IFE-1 versus IFE-3 pulldowns. Purple and green dots represent RNAs with >2-fold or <2-fold enrichment, respectively, and p<0.05. The majority of purple dots signify sequences of unknown function in WormBase, with the exception of abundant housekeeping mRNAs (see Supplemental Figure 1G, H). Blue-hatched boxes are shown as insets in E-G. Black dots identify germline translationally controlled mRNAs of interest initially identified in G as highly enriched by IFE-3 but not IFE-1. Also labeled are control tubulin (*tbb-2*) and GAPDH (*gpd-3*) mRNAs. H, I. RIP-Seq data was plotted relative to the measure of translational dependence on the IFE-1 isoform (fold mean R value, FMR; light blue dots) as measured by polysome microarrays in (41). Higher FMR values represent greater translational efficiency in wild type versus worms lacking IFE-1. Translationally controlled germ cell mRNAs previously shown to be activated by IFE-1 (red dots) or translationally repressed by IFE-3 (green dots) or unaffected by either IFE (black) are labeled (See refs (40,41,44). J. Quantification of binding of mRNA candidates to GFP::IFE-3 or mKate2::IFE-1 in RIP and qRT-PCR for validation of IFE-3 preferential binding. Cq values relative to total RNA samples and *tbb-2* control mRNA are plotted.

Because of the disparate IFE roles, we mined the RIP-Seq for candidate mRNAs substantially enriched by IFE-3 that contribute to those assigned phenotypic and translational functions. Germline mRNAs most evident by differential binding to IFE-3 were *spn-4, nos-1, pos-1, meg-1, mex-3, glp-1* and *oma-2* (Fig. 4E-G). By comparing their relative positions in the magnified volcano plots of IFE-1 vs total RNA, IFE-3 vs total RNA, and the IFE-1 vs IFE-3 plot, we could estimate which IFE mRNP binds these mRNAs stably. The RIP-Seq data shows that all 7 mRNAs were substantially enriched in the IFE-3 mRNP (Fig. 4F,G). The control mRNAs tubulin (*tbb-2*) and GAPDH (*gpd-3*) are not regulated by either IFE in the germline (40,41) and both were excluded from IFE-1 and IFE-3 mRNPs. Interestingly, *spn-4, nos-1, pos-1, meg-1, glp-1* and *oma-2* mRNAs were not enriched by IFE-1, but were also not excluded like *tbb-2* and *vpr-1* mRNAs (Fig. 4E). This behavior is consistent with dynamic and transient interaction of IFE-1 during positive translation initiation on these mRNAs as we previously demonstrated for this eIF4E isoform (41,44) see Discussion).

As a follow up to the newly identified mRNAs, we were interested in how mRNAs translationally regulated by either eIF4E might partition in the IFE-1 and IFE-3 mRNPs. Figures 4H and 4I plot the mRNA enrichment by each IFE relative to their translational dependence on IFE-1 (retrieved from translational microarrays in (41), rather than their p value. The fold mean R value (FMR) score is a ratio of translational efficiency in wildtype worms versus those with the *ife-1*(bn127) null mutation, derived from the earlier polysomal microarray study (41). mRNAs with higher FMR values are proportionally more dependent on IFE-1 for translation. We explored the catalog of all mRNAs previously shown to be translationally activated by IFE-1 (e.g. *lin-26, pos-1, glp-1, gld-1, mex-1, oma-1,* etc., red points; (41,44) and those translationally repressed by IFE-3 (*fog-1, daz-1, fem-3, etc.,* green points (40). These mRNAs were largely excluded from the IFE-1 mRNP (Fig. 4J, left side of plot). By contrast, most of these mRNAs were substantially enriched in the IFE-3 mRNP (Fig. 4K, right side of plot). Notable exceptions were *vpr-1, rab-7,* and *rab-5.* In addition, non-regulated mRNAs such as tubulin (*tbb-2*) and GAPDH (*gpd-3*) were equally positioned in graphs indicating they are excluded from both IFE-1 and IFE-3 mRNPs (Fig. 4E-G). Therefore, mRNP enrichment data for many cargo mRNAs (*lin-26, pos-1, glp-1, mex-1, oma-1, fog-1, daz-1, fem-3)* correlates with the repression of translation by IFE-3 and/or translational activation by IFE-1. Given their positions on the germ granule, our findings are consistent with IFE-1 receiving these mRNAs from IFE-3 at the time when they sorted and then recruited for translation (see Discussion). We validated the RIP-Seq data for *spn-4, nos-1, pos-1, meg-1, glp-1, oma-2* and *vpr-1* by direct qRT-PCR on IFE-1 and IFE-3 pull downs from the dual expressing strain. Both *spn-4* and *nos-1* mRNAs were enriched 12-20-fold relative to tubulin (*tbb-2*) mRNA levels (Fig. 4J). The *pos-1, meg-1, glp-1,* and *oma-2* mRNAs were enriched more than 5-fold, and even *vpr-1* was enriched 3-fold. Interestingly, IFE-1 also bound each of these mRNAs modestly, but none was enriched more than 2-fold (Fig. J). It is clear that both IFE mRNPs contain cargo mRNAs, but IFE-3 sequesters these regulated mRNAs far better than with IFE-1. These results show that select germline development mRNAs are being partitioned between the IFE mRNPs unevenly, likely leading to marked translational regulation by IFE-3::IFET-1 mediated repression. It is possible that mRNAs held in such translation factor-based repression is likely to restrict their synthesis (and hence function) to facilitate spatiotemporal development processes that are critical in germ cells and embryos.

### Repression of mRNAs by IFE-3 and IFET-1

Translational repression by the IFE-3 mRNP and translational activation by IFE-1 of the same mRNAs is a model of germ cell regulation to be tested *in vivo*. Reporter strains CRISPR/Cas9 engineered with in-frame GFP fusion to *mex-3, pos-1* and *spn-4* genes (a gracious gift of David Greenstein, Univ Minnesota (65) were evaluated for *in situ* expression upon loss of each IFE and its cognate 4EIP. The *mex-3, pos-1* and *spn-4* mRNAs favor binding to the IFE-3 mRNP complex (Fig. 4) and were assayed by *in vivo* fluorescence from GFP::MEX-1, GFP::POS-1 and SPN-4::GFP following depletion or absence of IFE-3, IFET-1, PGL-1/3 and IFE-1 (Fig. 5). Translational expression of these reporter mRNAs was assayed by fluorescence microscopy in the early germline (distal), during oogenesis (proximal) and post fertilization (embryo) by quantifying fluorescence intensities under defined, linear exposure conditions (see Methods).

**Figure 5:**
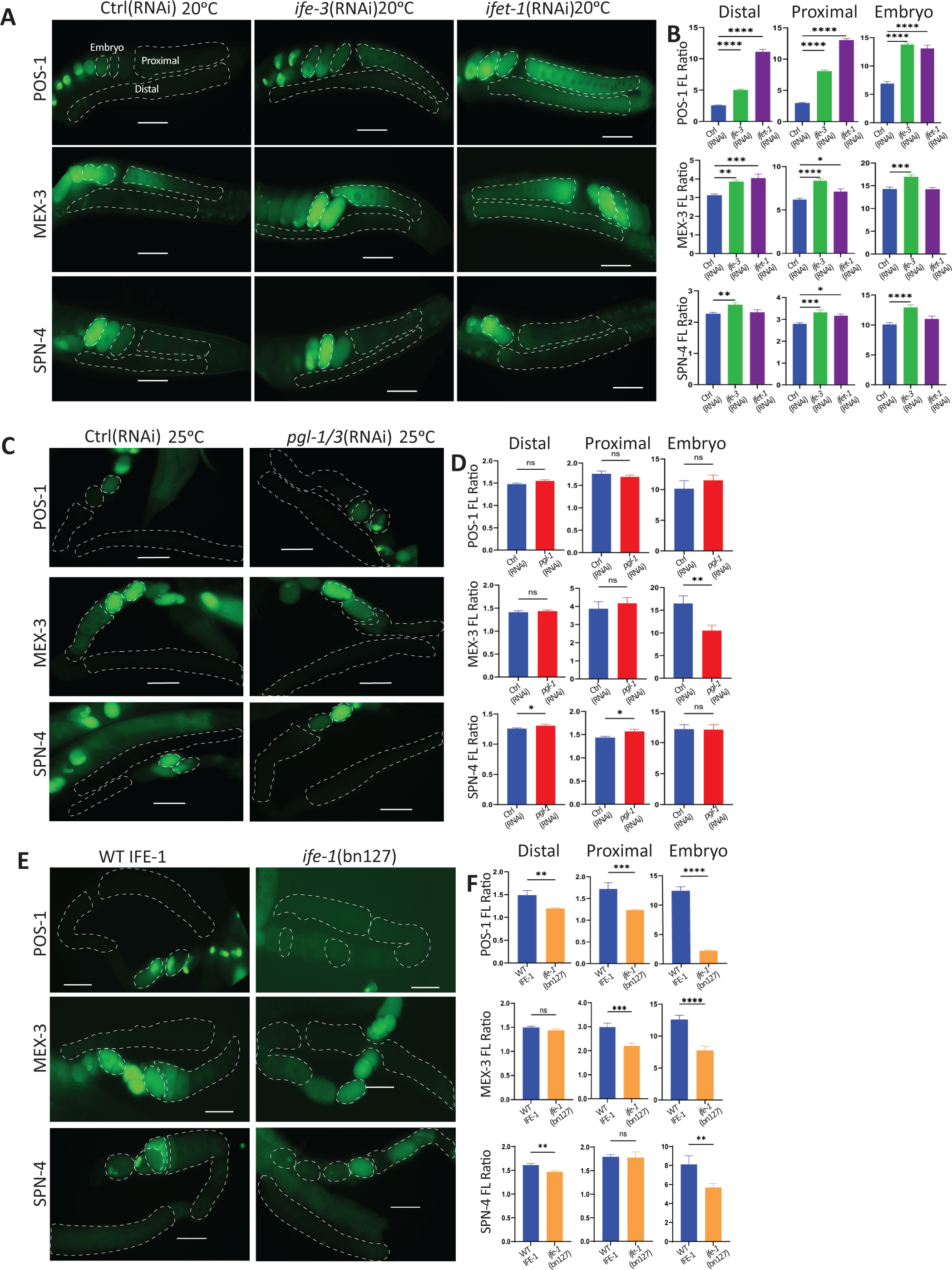
GFP fluorescent reporters for bound mRNAs show dependence on IFE-1, IFE-3 or their 4EIP partners. Expression of POS-1, MEX-3 and SPN-4 was studied following depletion or loss of *ife-1, ife-3, pgl-1/3 and ifet-1*. A. Expression each CRISPR/Cas9-fused reporter gene was imaged in the distal gonad, proximal gonad and uterine embryos following *ife-3*(RNAi), *ifet-1*(RNAi) or control (RNAi) treated worms at 20°C. B. Quantification of POS-1, MEX-3 and SPN-4 GFP fluorescence in the distal, proximal and embryos following *ife-3*(RNAi) and *ifet-1*(RNAi) at 20°C. Note the differing GFP intensity scales (y axis) between distal gonad and proximal gonad (maximally 2-5) versus embryos (maximally 10-20) reflecting higher native expression of these proteins by stage. C. Expression of POS-1, MEX-3 and SPN-4 upon *pgl-1/3*(RNAi) at 25°C; D. Quantification of POS-1, MEX-3 and SPN-4 GFP fluorescent in the distal gonad, proximal gonad and the embryo in *pgl-1/3*(RNAi) at 25°C; E. Expression of POS-1, MEX-3 and SPN-4 in *ife-1*(bn127) null mutant worms. F. Quantification of POS-1, MEX-3 and SPN-4 GFP fluorescent in the distal gonad, proximal gonad and the embryo *ife-1*(bn127) at 20°C. White hatched lines show the region of interest (ROI) used for fluorescence measurements. For all quantifications (n>15) native GFP fluorescence was normalized to an adjacent background signal in the same image and restricted to the linear response range of the camera (see Methods). Error bars are SEM. Statistical evaluation in B by ANOVA, in D and F unpaired t test. Designations: ns, not significant; ****, p <0.0001;***, p <0.001; **, p <0.01; *, p <0.05. Scale bars: 20µm

Loss of IFE-3 resulted in a strong enhancement of POS-1 expression (2.0 to 2.7-fold) in the distal and proximal regions of the germline as well as in uterine embryos (Fig. 5A, B). The de-repression of *pos-1* mRNA was even more pronounced following depletion of IFET-1, the 4EIP binding partner of IFE-3, in both the distal and proximal gonad (each 4.4-fold, Fig. 5B). Even in embryos, where *pos-1* mRNA becomes translationally activated only in posterior blastomeres (66), IFE-3 or IFET-1 depletion accounted for 2-fold de-repression. Most striking was that the *ife-3*(RNAi) and *ifet-1*(RNAi) embryos showed ectopic POS-1::GFP expression chiefly in the anterior blastomeres, leading to nearly uniform expression (Fig. 5A). There was substantial (but lesser) de-repression of *mex-3* and *spn-4* mRNAs upon *ife-3*(RNAi) and *ifet-1*(RNAi). Depletion of IFE-3 enhanced MEX-3 and SPN-4 expression reproducibly (up to 29%) in all regions of the germline and embryos (Fig. 5B), but IFET-1 depletion had a little effect except for some (29%) de-repression of *mex-3* in the distal gonad. The data indicate that all these mRNAs are held in a repressive IFE-3 mRNP complex, consistent with both IP-MS and RIP-Seq findings (Fig. 3, Fig. 4). Translational repression of *mex-3* mRNA by IFE-3 was previously observed by the Ryder lab, based upon a 3’ UTR binding site (67,68). We similarly demonstrated IFE-3-mediated translational repression using polysome resolution of *fem-3, fog-1, fbf-1, gld-1* and *daz-1* mRNAs (40). The current study confirms *mex-3* mRNA repression by the IFE-3 mRNP repression and extends it to *pos-1* and *spn-4* mRNAs. However, the severity of IFE-3 repression, and the contribution of IFET-1 to that repression, varies both among messages and in different regions of the gonad. Such findings indicate broad roles for this germline enriched eIF4E isoform in repressive mRNPs.

### Activation of repressed mRNAs by IFE-1, but no translational role for PGL-1

By contrast to IFE-3, the IFE-1 isoform binds strongly to IFG-1, the eIF4G ortholog in worms, indicating a substantial proportion of this eIF4E is engaged in positive translation initiation. PGL-1 is the direct 4EIP binding partner for IFE-1 (46,51);Kawasaki, 1998 #70} and is presumed to compete for binding to eIF4G. We addressed whether IFE-1 exerts positive or negative regulation on POS-1, MEX-3 and SPN-4 expression, and if PGL-1 plays any analogous role to IFET-1. Depletion of PGL-1 was shown to be highly effective at 25C (51,69) and as judged here by PGL-1::GFP reduction (Suppl Fig. 3). Importantly, loss of PGL-1 displaced IFE-1 from P granules (Fig. 2 and (40,46), but caused no significant decrease in mKate2::IFE-1 levels (Suppl Fig. 3). The loss of PGL-1 showed no significant effect on the expression of the three germline mRNA reporters (Fig 5C,D), except for a subtle increase of SPN-4 in the germline. The former might be explained by PGL-1’s role in sequestering a portion of IFE-1 (freeing the P granule fraction to find IFG-1). Unexpectedly, loss of PGL-1 substantially suppressed MEX-3 expression, but only in embryos. It is not clear why PGL-1 affects the *mex-3* mRNA only. Generally it appears that PGL-1 has no functional role in translational control of these mRNAs.

To assess IFE-1’s role directly, we first attempted *ife-1*(RNAi). Using the depletion of mKate2::IFE-1 as a read out, we found that depletion of IFE-1 was incomplete and showed no *ife-1* phenotype under the conditions we tested (data not shown). Instead, we crossed *ife-1*(bn127) into the POS-1, MEX-3 and SPN-4 reporter lines to generate homozygous null reporter strains. At the fertilization-permissive temperature of 20C (44,46,70), the lack of IFE-1 greatly decreased the expression of POS-1, MEX-3, and SPN-4, particularly in embryos where each of these mRNAs is best translated. In fact, POS-1 expression was suppressed by 82% in embryos lacking IFE-1, consistent with previous studies showing *pos-1* mRNA recruitment to polysomes depends on IFE-1 (44). Even the low residual expression of POS-1 in germ cells and oocytes (note the X-axis scale for Distal and Proximal graphs in Fig. 5F) was dependent on IFE-1 (20-28% lower). The contribution of IFE-1 to MEX-3 expression is substantial in late oogenesis and embryos (26% and 38%, respectively) but less robust than for POS-1. Interestingly, SPN-4 expression was responsive to IFE-1 only in embryos (30%; Fig. 5F). Given that POS-1, MEX-3 and SPN-4 natively show low translational expression throughout most of oogenesis, but become highly expressed at −1 or in cleaving embryos, these findings reinforce the primary role for IFE-1 in translational activation of regulated maternal mRNAs during oocyte to embryo transition (44, Friday, 2015 #1017,65). Notably, loss of individual eIF4Es or 4EIPs resulted altered localized translational expression patterns for these mRNAs that varied both up and down.

### IFE-1 and IFE-3 localize individually atop the coronal plane of the germ granule

Many lines of evidence have pointed to the importance of both functional and spatial juxtaposition of germline eIF4Es, their 4EIPs, other RBPs and helicases in germ granules. Multiple sub-compartments of germ granule have been reported in *C. elegans*, including Z granules, mutator foci, SIMR foci, and a P body structure (34,36,71). In light of our previous discovery that IFE-1 and IFE-3 occupy separate but adjacent positions in germ granules (40), we employed the same CRISPR/Cas9-constructed fluorescent fusions to carefully localize them relative to other important RNA regulatory proteins of the granule. Biochemical, genetic and *in vivo* microscopy evidence from this study confirm their distinct substructures, mRNA cargo, and functional activities. Others reported that IFE-3 is isolated as part of the PETISCO complex that functions in siRNA processing (61). Initially we hypothesized that IFE-3 may reside in Z granules because of its being part of the PETISCO complex (40). We subsequently crossed GFP::IFE-3 into a strain containing the definitive Z granule marker, ZNFX::mCherry. However, careful confocal imaging shows that IFE-3 and ZNFX-1 are non-overlapping in randomly positioned foci along the nuclear periphery (Fig. 6A, r=0.15). By contrast another known Z granule protein, LOTR-1 (45) is fully overlapping with ZNFX-1 (Fig. 6C, r=0.6). These findings make it clear that IFE-3 is not a resident of Z granule. Notably, the identical imaging strategy confirms our previous observation that IFE-3 localizes consistently adjacent to (partially overlapping) IFE-1 (Fig. 6B). IFE-1 has been reported to be resident in P granules proper because of direct protein-protein binding to PGL-1 (46,60). Given that IFE-3’s direct binding partner IFET-1 also associates with PGL-1, it seems that it could be a P granule resident. To address the positioning of the IFEs relative to other germ granule proteins, we create strains co-expressing PGL-1::GFP with mKate2::IFE-1 (Fig. 6E), GLH-1::GFP with mKate2::IFE-1 (Fig. 6F), and GLH-1::mScarlett with GFP::IFE-1 (Fig. 6G). PGL-1 binds directly to both IFE-1 and GLH-1 and has been reported to be more fluid in the germ granule compared to other granule residents (32,46,52,60). GLH-1 is the Vasa ortholog in *C. elegans* and is positioned more closely to the germ granule core (34,45). As expected, PGL-1 and IFE-1 colocalize and are fully overlapping, whereas IFE-3 and IFE-1 do not (Fig. 6E vs 6D). GLH-1 and IFE-1 are largely overlapping, but mKate2::IFE-1 red fluorescence appeared slightly external to GLH-1::GFP relative to the nuclear membrane, even by conventional fluorescence microscopy (Fig. 6F, enlarged last panel). Any trend toward separation was surprising because only PGL-1 separates IFE-1 and GLH-1 in protein-protein interactions. Finally, IFE-3 was found to be non-overlapping with (and always external to) GLH-1 in a manner more pronounced than IFE-1 (Fig. 6G vs 6F, enlarged last panels). The hints of a stratified and oriented structure involving IFE-1, IFE-3, PGL-1 and GLH-1 prompted the need for localization of these germ granule proteins using high resolution confocal Airyscan microscopy (Fig. 6H-K). IFE-1 and IFE-3 localized separately with IFE-3 puncta occasionally forming a halo around IFE-1 loci (Fig. 6H). PGL-1 was seen to be a broader, flattened almost triangular structure into which IFE-1 intercalated as smaller puncta (Fig. 6I). When co-expressed with GLH-1, IFE-1 was now visible as multiple (2–4) red puncta mostly apical to the GLH-1 raft-like disc along the nuclear periphery (Fig. 6J). Finally, IFE-3 routinely formed two lateral puncta at the ends of the red fluorescent GLH-1 disc, which was again more closely adjacent to the nucleus (Fig. 6K). Taken together, fluorescent imaging within the condensate shows a stratified arrangement of proteins. GLH-1 and PGL-1 inhabit a core on which the IFEs are non-randomly localized foci over the corona (Fig. 6L).

**Figure 6:**
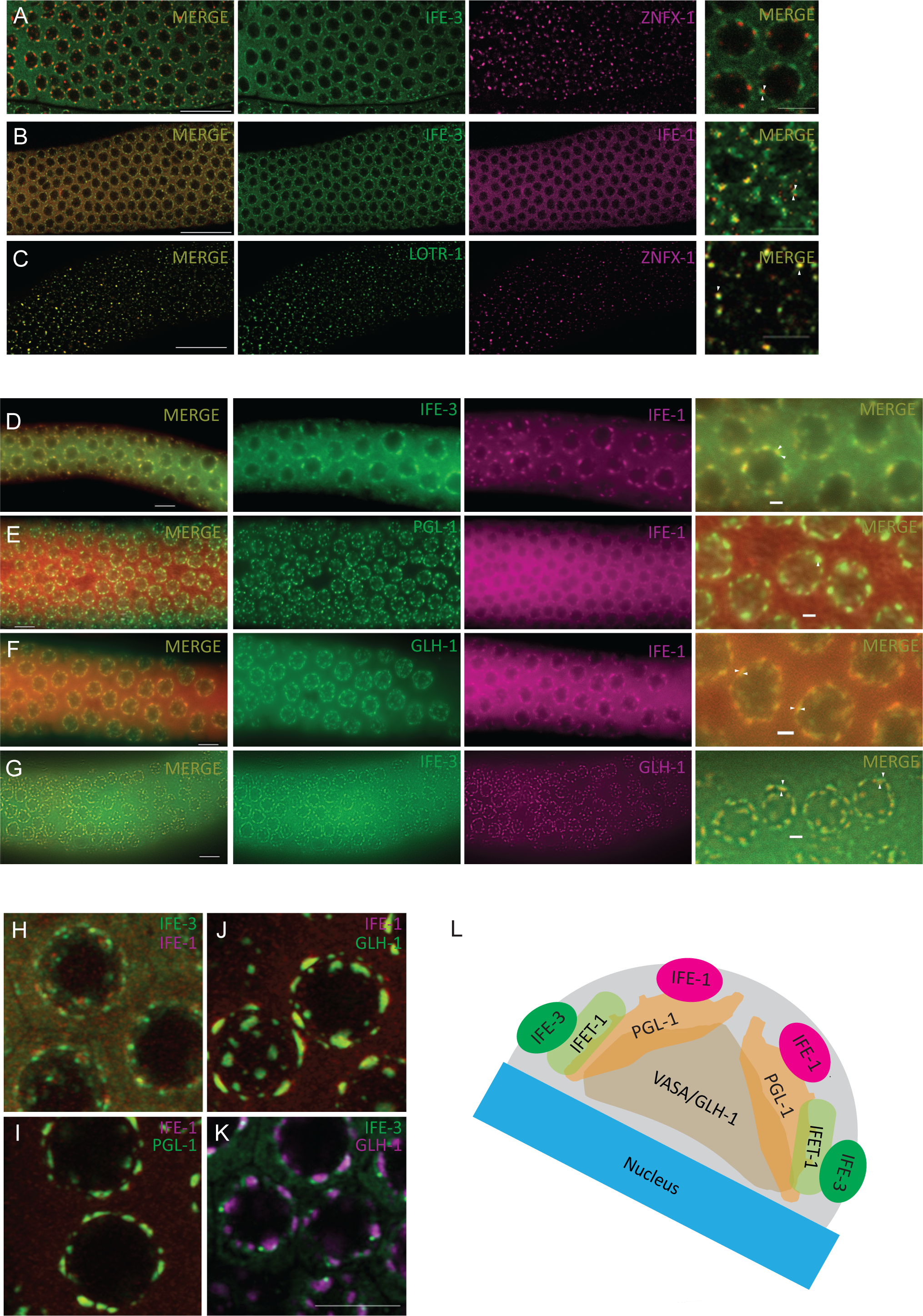
Separated and outward location of IFEs in the germ granule correlates with their functional interface with translation apparatus. Dissected gonads were subjected to high resolution and confocal microscopy to localize fluorescent fusions of canonical P and Z granule proteins. High magnification merged images are to the right. A. Comparison of IFE-3 to ZNFX (Z granule). GFP::IFE-3 does not colocalize with ZNFX-1::mCherry, a marker for Z granules. (correlation regression r=0.15) B. GFP::IFE-3 localizes adjacent to mKate2::IFE-1 in germ granules under the same conditions (correlation regression r=0.6). C. LOTR-1::GFP and ZNFX-1::mCherry colocalize in Z granules as expected (correlation regression r=0.6). D. mKate2::IFE-1 localization adjacent to GFP::IFE-3 as imaged by standard fluorescence microscopy. E. Complete overlap of mKate2::IFE-1 with its 4EIP PGL-1::GFP; almost no PGL-1:GFP is in the soluble portion, whereas substantial mKate2::IFE-1 is soluble. F. mKate2::IFE-1 is slightly separated from GLH-1::GFP and lays more external in the peripheral ring around the nucleus. G. GFP::IFE-3 is significantly separated from GLH-1::mScarlett and is also more external to the nucleus. H. mKate2::IFE-1 and GFP::IFE-3 are visible as clearly separate puncta along the outer layer of germ granules by super-resolution Aryscan fluorescence microscopy of fluorescently tagged germ granule proteins. I. mKate2::IFE-1 forms tight puncta on the surface of a broader PGL-1::GFP that is now resolved by super resolution. J. mKate2::IFE-1 tight puncta are now further separated from an even broader GLH-1::GFP raft-like structure. K. GFP::IFE-3 spots are also separable from GLH-1::mScarlett but are not as tight and appear to localize more laterally on the GLH-1::mScarlett raft structure. (A-C. 63x standard confocal microscopy, scalebar 20µm, D-G 100x oil standard fluorescence microscopy, scale bar 20µm and 5µm. H-K. Airy Scan confocal microscopy, scalebar 5µm.) L. Model of germ granule proteins in a stratified arrangement with IFE-1 and IFE-3 separated near the surface, and PGL-1 and GLH-1 more internal in the structure. Unlike most germ granule components, both eIF4E isoforms both have an observable presence in the soluble germ cell cytoplasm.

## Discussion

### Translational repression or initiation by two different eIF4Es

Involvement of the mRNA cap-binding protein eIF4E in both repression and translation initiation was discovered in *Xenopus* oocyte meiotic maturation over 23 years ago (72). A 4EIP partner called Maskin was found to mediate repression by preventing association of eIF4E with eIF4G to form the eIF4F complex for mRNA translation initiation (8,73–75). Hormone-induced phosphorylation of CPEB caused dissociation of the eIF4E from the Maskin mRNP to facilitate poly(A) tail elongation and binding to a positive translation initiation complex (76). Here we propose an alternative model in *C. elegans* oocytes and embryos that utilizes two eIF4E isoforms to mediate repression and activation in the germ granule: IFE-1 and IFE-3. Each has a dedicated 4EIP that hold them in the germ granule. IFE-1 binds PGL-1 and IFE-3 binds IFET-1 (40,44,46,48). The current study indicates that one eIF4E:4EIP acts on controlled mRNAs in a repressed mRNP (IFE-3:IFET-1) while the other mediates translational activation (IFE-1:PGL-1 moving to IFE-1:IFG-1). Activation leads to initiation via IFG-1 (eIF4G) binding and recruitment to ribosomes outside of the germ granule.

This model (Fig. 7) is most consistent with our various observations both *in vitro* and *in vivo*. We isolated the stable IFE-3 mRNP to characterize its protein composition and cargo mRNAs led to the conclusion that its primary role may be sequestration of germ granule mRNAs, with numerous repressor RBPs associated and the *pos-1, mex-3* and *spn-4* mRNAs among the most enriched (Fig. 3 and 4). Depletion of either IFE-3 or its binding partner IFET-1 improved synthesis of POS-1, MEX-3 and SPN-4 GFP fusions *in vivo*. It is rather striking that RNAi against the canonical eIF4E isoform would cause enhanced translation of *pos-1, mex-3* and *spn-4* mRNAs over wild type levels. Interestingly, the amount and place of activation was not the same for all the reporters in germline or embryos. Depletion of the other germ granule eIF4E (IFE-1) had the opposite effect on expression POS-1, MEX-3 and SPN-4, indicating it was necessary for recruitment of these controlled mRNAs to ribosomes. Clearly there are translational control checkpoints involving IFE-1 and IFE-3 that govern when and where certain mRNAs becomes translated. The simplest explanation is that germ granule transcripts are released from the IFE-3 mRNP complex to increase their availability to IFE-1 to translate them (see below). Their unique translational regulation in various stages of germ cell development and in early embryos is critical for developmental progression to mature gametes and viable embryos. Loss of POS-1 results in maternal-effect embryonic lethality caused by abnormal cell-fates and cell divisions because POS-1 regulates *apx*-1 mRNA translation (77). POS-1 also represses maternal *glp-1* mRNA translation in embryonic posterior blastomeres at the same time that SPN-4 activates translation of *glp*-1 mRNA in anterior blastomeres (66). MEX-3 is a KH-type RBP that controls maternal *pal-1* mRNA translation (78). Ironically, we previously showed that *pos-1, pal-1* and *glp-1* mRNAs all rely on the IFE-1 eIF4E isoform for efficient translational activation (41,44).

**Figure 7:**
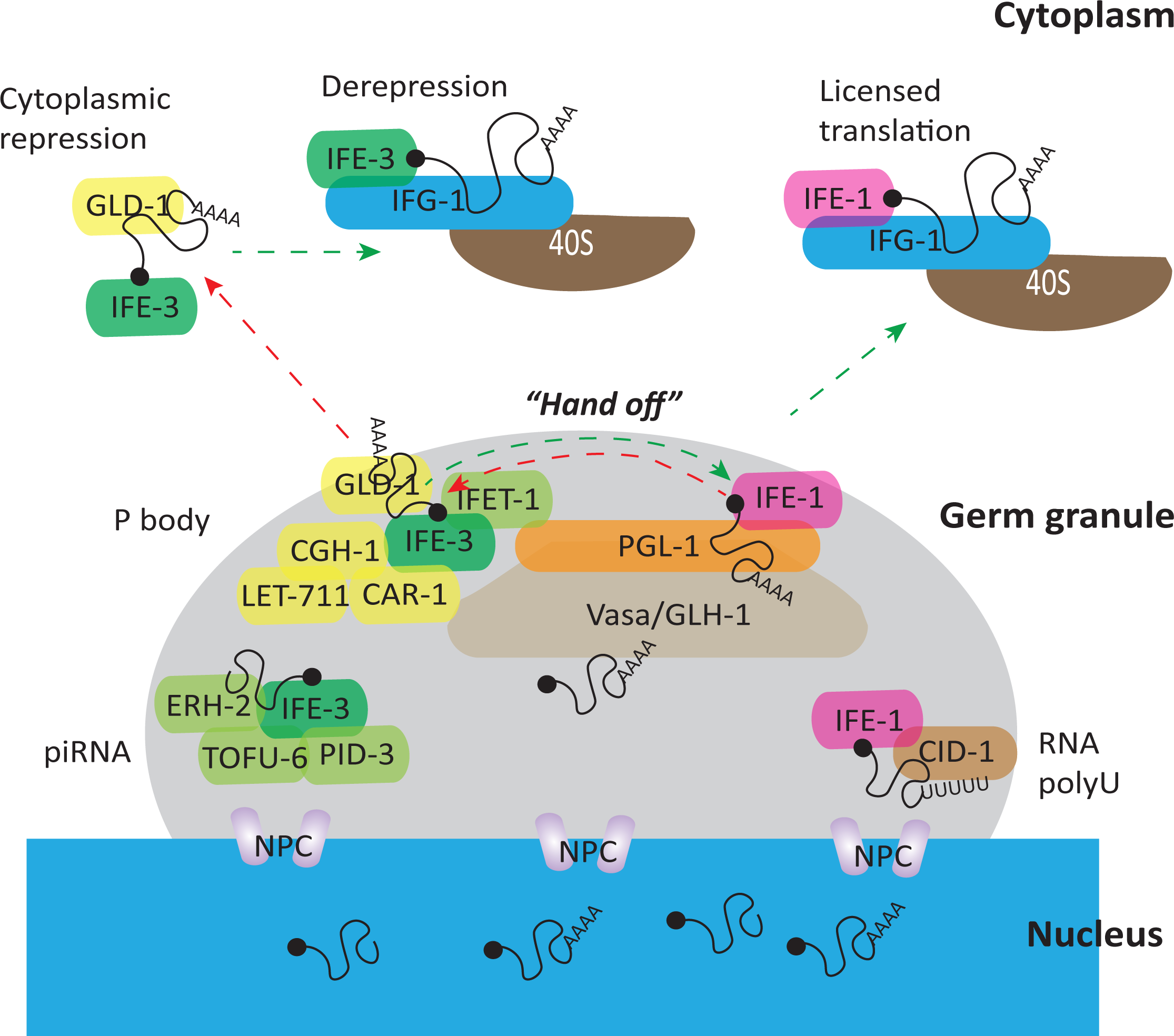
Multiple RNA functions of the cap-binding IFEs residing in germ granules. Conceptual model of germ granule compartmentalization that involves multiple IFE-1 and IFE-3 mRNP complexes for various RNA cap-binding roles in 1) piRNA processing, 2) P body function, 3) mRNA export for translation, and 4) RNA polyuridinylation. The stratified arrangement with IFE-1 and IFE-3 located near the “outside” of the germ granule is suggested by their morphology as adjacent puncta localized outward from the nucleus relative to PGL-1 and GLH-1. It is further supported by the opposite functions implied by their protein binding partners and the mRNAs that they retain or translationally regulate. The PGL-1::GLH-1 scaffold/raft structure coordinates both IFE mRNPs through interactions with PGL-1. IFE-1 binds directly and IFE-3 indirectly through the 4E-T ortholog IFET-1. P body-associated and RBP repressor proteins (CGH-1, CAR-1, LET-711, GLD-1) bound to IFE-3 may hold some mRNAs in repression within the germ granule. When appropriate, those mRNAs are handed off to IFE-1 along PGL-1 for more immediate access to eIF4G (IFG-1), resulting in active translation initiation in the cytoplasm. In this study we identified germline mRNAs that are activated by IFE-1 and stably bound/repressed by IFE-3 (Fig. 4) as evidence for this mechanism. The “inward to outward” stratification puts Vasa\GLH-1 in a position to receive RNAs transiting through the germ granule and perhaps exercise the “licensing” role proposed by others. GLH-1 coordinates small RNA assembly on or off of mRNAs via the Argonauts/CSR-1 as a way to determine their translational fate. IFE-3 mRNPs are also likely released from the germ granule for mRNAs carrying a sequence-specific RBP translational repressor like GLD-1. These would eventually be resolved in the cytoplasm allowing IFE-3 to act as a positive eIF4E and initiate translation through IFG-1. IFE-1 and IFE-3 may also have secondary cap-binding roles in the germ granule involving non-mRNAs. IFE-3’s role in piRNA processing in the PETISCO complex is already established. IFE-1 association with the poly(U) polymerase CID-1 implies that it may coordinate the caps of RNAs being marked for degradation. However, no further evidence has been discovered for a role in polyuridinylation.

There are now several confirmations that IFE-3:IFET-1 is an important translational repressor heterodimer in the germline. We demonstrated by polysome fractionation that loss of either IFE-3 or IFET-1 de-repress translation of *fem-3, fog-1, fbf-1, gld-1* and *daz-1* mRNAs (40). Independently, others have demonstrated that IFE-3 directly mediates repression for *mex-3* mRNA (67). These critical germline mRNAs and others are sequestered by IFE-3 mRNPs, but are largely excluded from IFE-1 mRNPs (Fig. 4). Coincidentally, these same mRNAs were also shown to require IFE-1 for efficient translation (41). IFE-mediated sequestration does not occur on all germline transcripts since *tbb-2, gpd-3* and *vpr-1* mRNAs were excluded from both IFE-1 and IFE-3 isolations (Fig 4E, F, G). It is worth noting that *mex-1* mRNA was shown to require IFE-1, but not IFE-3, for efficient translation, and is not repressed by IFE-3 either (40,44). Yet *mex-1* mRNA transits through germ granules and is translationally regulated in late oogenesis/embryos (79,80). This suggests that other modes of translational repression (e.g. GLD-1, GLP-1 in the germline or LIN-41/OMA-1/2 during the oocyte to embryo transition) may be independent mRNA repression complexes that can also funnel into IFE-1-mediated translation initiation (81,82).

By contrast, all studies to date show that IFE-1 supports only positive translational eIF4E roles (29,33,41,44,46,70). Absence of IFE-1 [*ife-1*(bn127)] caused substantially reduced (but not abolished) expression of POS-1, MEX-3 and SPN-4, particularly in the oocyte to embryo transition. Other evidence shows they are members of substantial subset of germline/embryo mRNAs that uniquely require this eIF4E isoform for efficient translation (*pos-1, vab-1, glp-1, oma-1, lin-26, let-49, mex-1, rab-5, −7*, etc., (41,44). Yet most of these mRNAs were not isolated in IFE-1 mRNPs but rather in IFE-3 mRNPs, suggesting that only repressed complexes are stable to such purification. Our findings show that individual eIF4Es or 4EIPs direct a spectrum of localized translation patterns for *pos-1, mex-3*, *spn-4*, and other germline mRNAs based on germline region and mRNA translational character. The data suggests that eIF4Es and 4EIPs act in a combinatorial fashion to influence de-repression and ribosomal recruitment during the course of development.

### The case for mRNA hand off between germ granule eIF4E isoforms

It was advantageous to be able to characterize both the proteins and mRNAs that reside in the IFE-1 and IFE-3 mRNPs, and compare them to one another. In using both proteomic and RIP-Seq methods, we could conclude that we captured the most prevalent forms of each complex and were intrigued to see how divergent the protein and RNA species were for two similar germ cell eIF4Es. However, our interpretation is subject to several caveats: 1) mRNPs formed around the IFEs are likely of variable stability such that the most dynamic forms may be less stable and therefore underrepresented by affinity pulldowns. This is especially true of translation initiation and ribosomal complexes where associations are transient and catalytic in nature. Translational recruitment involves unstable, dynamic interactions of the mRNA with eIF4E/eIF4G, eIF4A, eIF3 with the 40S subunit only during initiation; they are released upon joining of 60S (4,7,83). These initiation intermediates do not remain bound in the absence of catalytic inhibitors (5,7,8,43,63). 2) The protein and mRNA species identified do not represent a single complex; more likely they represent a composite of several co-existing complexes. The clearest example is strong detection of both IFE-1:PGL-1 enrichment, showing the mRNP form found perhaps exclusively in the germ granule, and IFE-1:IFG-1, showing an eIF4F mRNP that is expected to exist only in soluble cytoplasm. 3) Purifications from whole worms are also composites of diverse cell types and stages. Both IFE-1 and IFE-3 are much more strongly expressed in the germline than in soma (40), and consistent with our identification of primarily proteins and RNAs that are also germline expressed. But even mRNPs formed in mitotic germline stem cells and early meiotic cells could be extensively remodeled by the time they differentiate into oocytes, sperm and ultimately embryos.

There is extensive evidence that germ cell eIF4Es make discrete mRNPs in many plant and animal species (reviewed in (4,12,27,30,72). The nematode IFE-1 and IFE-3 mRNPs each contain unique proteins and transcript pools in separable sub-compartments in the germ granule. Current literature suggests the biophysical nature of protein and RNA components within germ granule condensates dictate the substructure fluidity, which adds an extra layer of potential organization to mRNA pathway (32,33). Given that translationally active eIF4Es are not expected to stably bind mRNAs, we learned more than expected from the RIP-Seq characterization of IFE-1 and IFE-3 mRNPs. In particular, there was a consistent pattern for the partitioning of their mRNA cargos. IFE-3 sequestered nearly all of the mRNAs that have been shown to be positively regulated by IFE-1 (*lin-26, vab-1, glp-1, gld-1, oma-1, nos-1, pos-1, let-49).* These mRNAs are excluded from IFE-1 mRNPs (41, Henderson, 2009 #824), Fig. 4H, I) as would be expected of translation initiation complexes. Likewise, a correlation with IFE-1 translational activity was seen for mRNAs shown to be translationally repressed by IFE-3 (*fog-1, fem-3, daz-1, tra-2*), again not substantially bound by IFE-1 mRNP. From these data we infer that these regulated mRNAs are handed off from one IFE to the other during a switch from repression to active translation (and perhaps vice versa). IFE-3 was found in complex with proteins from several small RNA pathways, so it is likely that Argonautes present there might cooperate in loading small RNAs onto cargo mRNAs. The miRNAs may license mRNAs for either continued repression via IFE-3:IFET-1, or to be handed off to IFE-1:PGL-1, potentially to shuttle them out of germ granules to IFG-1 for translation (Fig. 7). We noted mRNAs that were exceptions because they were excluded from both mRNPs. The *tbb-2* and *gpd-3* mRNAs (not translationally regulated in germ cells) and a group of strongly IFE-1-dependent mRNAs (*vpr-1, grl-20, rab-5, rab-7, ran-1*) that have no critical role in germline development may circumvent germ granule mediated licensing. The model (Fig. 7) is an attempt to explain how mRNAs travel across the germ granule, getting progressively decorated or unwound by RNA helicases, eIF4Es, 4EIPs, miRNAs, Argonautes and RBPs and ultimately exit the germ granule condensates to bind ribosomes.

Interestingly, loss of PGL-1 neither enhanced nor repressed the expression of POS-1 or SPN-4 despite almost complete depletion (Fig. 5 and Suppl Fig. 3B). MEX-3 expression was diminished in embryos only when PGL-1 was depleted, for reasons that are unclear. However, the results clearly show that the IFE-1-specific 4EIP is not acting as a translational repressor, in contrast to the role of IFET-1. PGL-1 binds to IFE-1 using a similar motif to IFG-1 (eIF4G) and was assumed to compete with cap-dependent translation (46,84), an assumption that was never functionally substantiated. Our findings indicate that PGL-1 plays no direct role in translational control of *pos-1, mex-3* and *spn-4* mRNAs. IFG-1 and PGL-1 were among the top three proteins coprecipitated with IFE-1 (Fig. 3). We suggest that they represent two distinct IFE-1 mRNPs because they bind at the same site on this small eIF4E protein. But the findings also suggest that IFE-1 dissociation from PGL-1 is not the rate-limiting step in active ribosomal recruitment of IFE-1-dependent mRNAs *in vivo*. Others previously showed that artificial tethering of PGL-1 to mRNAs prevented their translation (85). Consistent with both notions is a model where the native function of the PGL-1 4EIP is to mediates germ granule association of both IFE-1 and IFE-3 mRNPs. By concentrating eIF4Es at the external face of germ granules, PGL-1 could promote an mRNA swap, then release the IFE-1 mRNP to IFG-1 for cap-dependent translation initiation (Fig. 7). In unique instances, PGLs may even facilitate the recruitment to IFG-1 (explaining diminished MEX-3 in *pgl-1/3*(RNAi) embryos). More intriguing is the interplay of IFE-1 and IFE-3 mRNPs because of the role of IFET-1, which is also reported to associate with PGL-1 (48,49). Taken together, our findings are consistent with a remodeling of both IFE-1 and IFE-3 mRNPs within native germ granules. The purpose of the remodeling is to achieve a functional “hand-off” of cargo mRNAs from IFE-3 to IFE-1 on the way to out of the granule (Fig. 7). Progressive remodeling forms an mRNP ready to interact with eIF4G to form the cap-dependent eIF4F (IFE-1:IFG-1 p170), and then addition of eIF3 and 40S ribosomal subunit for active initiation (3,5,12). The swapping of PGL-1 binding for IFG-1 binding may precede or be coincident with the IFE-1/IFE-3 remodeling. Because PGL-1 remains germ granule associated and eIF4G (IFG-1) remains soluble under non-stressed conditions (86) and data not shown) it may be that the two eIF4E binding proteins need not compete for binding due to compartmentalization.

### Remarkably different cap-binding mRNPs

It was not intuitive that the two germline eIF4Es would differ substantially in their protein-protein interactions and cargo mRNAs. IFE-1 and IFE-3 are 25 kDa proteins that share 48% sequence identity (and much higher sequence/structural similarity) and bind m^7^GTP with similar affinities (29,87). High sequence identity prevented isoform-specific immunoprecipitation until we separately tagged each *ife* gene using CRISPR/Cas9 genomic engineering (40). In this study mKate2::3xMyc::IFE-1 and GFP::3xFlag::IFE-3 proteins were separately affinity purified from a co-expressing worm strain and their protein/RNA complexes subjected to IP-MS and RIP-Seq to determine the protein composition and mRNA cargo of each cap-binding mRNP. To our surprise, the constitution of the IFE-1 and IFE-3 mRNPs differed almost completely (Fig. 3). IFE-1 was found complexed with a small pool of proteins largely involved in active translation initiation such as the eIF4G-1 ortholog IFG-1, PAB-1 and a few ribosomal proteins. This was notable because translation initiation interactions are transient, so that purification of catalytic complexes (e.g. with eIF3, eIF2, etc.) is unlikely in the absence specific inhibitors (2,5,7,8,75,88). An exception is the moderately stable eIF4F complex (eIF4E, eIF4G and in some species, eIF4A), which in our IP-MS findings appears chiefly as IFE-1 with IFG-1. We previously showed that the long isoform (p170) of IFG-1 promotes mRNA cap-dependent translation initiation that is essential to prevent ectopic germ cell apoptosis (55,56,89). Strong association of IFG-1 with one eIF4E isoform but not the other indicates that IFE-1 has a greater propensity to form a productive eIF4F translation initiation complex than IFE-3 (5,74). However, the most enriched protein to coprecipitate with IFE-1 was CID-1 (also called CDE-1, PUP-1) that encodes the ortholog of human TUT7 poly(U) transferase. CDE-1/CID-1 polymerase elongates 26S rRNA and CSR-1-bound small RNAs, marking them for degradation (58,59). CID-1 is necessary for meiotic chromatin methylation and compaction (57) likely in conjunction with the 22G RNA processing by CSR-1. (CSR-1 was modestly enriched in both IFE-1 and IFE-3 samples, but neither was statistically significant). It is unclear what relationship 3’ poly(U) elongation of other RNAs has to mRNA 5’ cap-binding. However, given IFE-1’s positive role in oocyte and sperm protein synthesis, a putative IFE-1/CID-1/CSR-1 complex may provide feedback signaling already in germ granules via miRNA regulation and ribosome biogenesis pathways. PGL-1 was of course also bound to IFE-1 as it is a cognate 4EIP that is known to have direct protein-protein interaction with only this eIF4E isoform (46). PGL-1 also binds directly to GLH-1, an intrinsically disordered domain protein with links to small RNA pathways (22,37), further links IFE-1 to miRNA regulation, albeit indirectly. PGL-1 could be sieving miRNA/Argonaute-bound mRNAs destined for translation or degradation with the help of helicase activity of GLH-1 and WAGO-1 (90). PGL-1 was also linked to PRG-1 function in the 21U RNA biogenesis pathway (91).

Genetically and biochemically, IFE-3 plays very different roles from IFE-1 in germline RNA regulation. Several studies demonstrate translational repression of mRNAs by IFE-3 (40,67). The translational repressor IFET-1 (ortholog of vertebrate eIF4E-T) was the second most enriched protein in the IFE-3 mRNP (Fig. 3). IFET-1/SPN2 was previously characterized as an isoform-specific 4EIP (48,92). Interestingly, only germline mRNAs appear subject to IFE-3:IFET-1 repression, and the IFE-3 mRNP-bound mRNAs characterized here are only expressed in germ cells. *C. elegans* IFE-3 is the direct ortholog of canonical human eIF4E-1 and has a presence in somatic tissues. Its repressive role may be lost during embryogenesis where it is the only essential eIF4E, ostensibly as the positive eIF4E in the new somatic cells (29,40,93) and data not shown). Outside of translation, IFE-3 was also implicated in small RNA pathways, particularly as part of the PETISCO complex by pulldown of other subunits (61). Indeed, all of the PETISCO proteins, TOFU-6, PID-3, ERH-2 and TOST-1, were among the most abundant partners to IFE-3 in our study (Fig. 3). PETISCO functions in 21U RNA (piRNA) biogenesis in germ cells and embryos. Defects in other PETISCO proteins likewise result in a gonad masculinization phenotype (62,94). IFE-3 is likely present for its affinity for m^7^GpppN caps, which are present on the 5’ of 21U precursor transcripts (61,95). Perhaps IFE-3 is a promiscuous cap-binding protein utilized broadly in RNA processing capacities. Proteins of the Argonaute miRNA pathway (ALG-1, −2, AIN-1, −2) were also found to be part of IFE-3 mRNPs. Perhaps most striking, remaining partners were well characterized translational repressor RBPs (GLD-1, RNP-8 and MXT-1) and P body/mRNA degradation proteins (CGH-1, CAR-1, LET-711; Fig. 3). Interestingly, proteins of the repressive LIN-41 mRNP were not found in the IFE-3 mRNP (except GLD-2) highlighting a diversity of 3’ UTR-binding complexes (65). These data suggest that IFE-3 resides in several, but not all, translationally repressed mRNA complexes. Other bound proteins link IFE-3 to potential positive translational regulation, including one subunit of eIF3 (eif-3.G) and the rpS6 kinase RSKS-1. S6 kinase plays a role in 40S ribosome subunit activity that modulates TOP mRNA translation (96) and the *C. elegans* ortholog RSKS-1 is critical for stem cell maintenance and longevity cues (97,98). IFE-3 also binds to the GLD-2, - 3 and GLS-1 germline proteins. The GLD-2/3 complex plays a well characterized role in cytoplasmic polyadenylation (99–103). GLS-1 is a novel P granule and polysome component that enhances the enzymatic activity of the GLD-4 poly(A) polymerase for germ cell mRNA activation (104,105). Together these implicate IFE-3 in various other positive translational roles. Most unusual, but significant, were proteins involved in small nuclear protein pathways (NOL-56, SNR-4, SNR-7, and SMN-1). Since we do not observe substantial GFP::IFE-3 in the nucleus at any stage, the importance of these interactions is unclear. Nuclear functions have been ascribed to mammalian eIF4E in mRNA 3’ end cleavage and export (106), but there is no current evidence from that IFE-3 plays such a role. Overall, the uniquely different mRNP composition for each IFE suggest they participate in different functional RNA processes during germ cell development and embryogenesis. Given that both exist in separable substructures of the germ granule, as well as free in the cytoplasm, each IFE may have multiple functions (that presuppose multiple mRNP types) in mRNA trafficking, polyadenylation, translation, and turnover.

Just two proteins were found robustly bound to both IFE-1 and IFE-3 (>70-fold enrichment, Fig. 3E) in roughly equal proportions. These were Mextli (MXT-1) and GIGYF2 (GYF-1), both known 4EIPs (19,53,54,107). GYF-1 was previously identified as the 4EIP for yet another *C. elegans* eIF4E, IFE-4, the nematode homolog of eIF4E2/4EHP (23,29). IFE-4:GYF-1 represses mRNA in *C. elegans* neurons. This complex was recently suggested to loop capped mRNAs that cannot be recruited to ribosomes due to damage or NMD targeting (23,53,107,108). Evidence points to GYF proteins exerting translational repression in all cells, irrespective of the eIF4E isoform partner. Mextli is another indiscriminate 4EIP initially identified in *Drosophila* and later in *C. elegans* that is a positive regulator of translation (54). It has been suggested to serve as a functional analog of eIF4G-1 because it contains an MIF4G domain and interacts with eIF3 (54). It is formally possible that MXT-1 could substitute for IFG-1 with IFEs to facilitate translation initiation. However, we and others have published extensive biochemical and genetic evidence that the *C. elegans* IFG-1 has essential functions in embryogenesis, larval development, miRNA regulation, prevention of germ cell apoptosis and somatic lifespan that are not suppressed by MXT-1 (55,56,89,109–111).

### What phenotype and localization tell us about germ granule life

Null mutations in the *ife-1* and *ife-3* genes cause defects in germ cells and embryos, but at different phases in their progression. Loss of IFE-1 causes a temperature-sensitive arrest in secondary spermatocyte budding and a temperature-insensitive reduction in oocyte growth (41,44,46). The overall effect on fertility is compounded because late-stage oocytes await a maturation signal from major sperm protein (MSP) to resume meiosis. MSPs were shown to be deficient in IFE-1-depleted worms (70), but polysome analysis showed it was due to lower *msp* mRNA levels rather than translational control (41). We also observed a morphological consequence of IFE-1 loss that may arise from reduced oocyte/sperm production. Broadening of hermaphrodite and male gonads was obvious upon dissection. Perhaps more relevant was a disruption of the gonad rachis lattice, or at least IFE-3 association with it. Whether lattice reorganization is due to direct interaction of IFE-1 with IFE-3, as observed in the IP-MS, is not known. But no similar displacement of IFE-3 from germ granules was observed (Fig. 1D). IFET-1 is the direct binding partner to IFE-3 and was required to hold IFE-3 in both rachis structures and germ granules (Fig. 2F and (40). This is further evidence of a hierarchy of interacting proteins that differs between these different locations. Regarding its own direct binding partners, IFE-1 was not required to hold either PGL-1 or GLH-1 in the germ granules (Fig. 1). We can infer that loss of IFE-1 does not substantially affect germ protein composition, suggesting that it is the most “outward-facing” protein in the structure (Fig. 7). Conversely, the depletion of IFE-3 did diminish IFE-1 in germ granules, but with no overall change in soluble IFE-1 fluorescence. This indicates that the IFE-1:IFE-3 interaction is of some consequence relative to their condensate localization. Complete absence of IFE-3 does not disturb the IFET-1 or GLH-1 localization to germ granules, suggesting that IFE-3, just like IFE-1, is not integral to the germ granule protein composition. We propose a model in which IFE-3 interaction with IFE-1 may serve to deliver previously repressed mRNAs to a preliminary initiation complex (Fig. 7). In that scenario, PGL-1:GLH-1 tethers both the IFE-3 mRNP and the IFE-1 mRNP in the germ granule. Once mRNA cargo is delivered to the IFE-1 mRNP, it exits the germ granule. In the cytoplasm it associates with IFG-1 (eIF4G) for recruitment to 40S ribosome via cap-dependent translation initiation (Fig. 7).

The IFE-1, −2, −3, 4 and −5 eIF4E isoforms were initially thought to have largely overlapping functions because their protein sequences/structures and cap-binding affinities were so similar. In addition, all *ife* null strains except *ife-3* are viability, which is unusual for protein synthesis factors with general required cellular functions (29,87). But the notion of functional redundancy was dispelled by the unique fertility and sensory phenotypes observed, and later the characterization of isoform-dependent mRNA subsets for four of the IFEs (29). IFE-1 recruits translationally repressed germ cell mRNAs such as *glp-1, oma-1, pos-1, mex-1, mex-3, vab-1,* etc. in late oogenesis to promote maturation (41), as well as unidentified mRNAs in secondary spermatocytes to promote budding (44,46). IFE-2 functions in both soma and germline; in the latter it translationally recruits *msh-4* and *msh-5* mRNAs to facilitate late meiotic crossover events and chiasmata integrity in oogenesis (42). Curiously, only *ife-3* null mutations cause embryonic lethality, but typically in the F2 generation after substantial germline sex determination defects in F1s (40,93). Translationally it is clear from our previous studies and figure 5 that IFE-3:IFET-1 mediates repression of the germline mRNAs *mex-3, fem-3, fog-1,* and *daz-1*, but IFE-3 modestly enhances translation of *tra-2* (40,67). IFE-4 is a somatic-only isoform expressed in muscle and neurons that translationally recruits *daf-12, egl-15* and *kin-29* mRNAs which are important for vulval innervation and egg-laying (43). Only IFE-5 remains uncharacterized; its gene likely derives from a recent gene duplication of *ife-1*, but is expressed at much lower levels in the germline and its loss is of no phenotypic consequence (29). These studies confirm that all 5 eIF4Es are unequal in their roles for mRNA translational control, consistent with the unique subsets of IFE-dependent mRNAs identified by our many polysome profile experiments on worms deficient in each eIF4E isoform (40–44).

There are intriguing outstanding questions about the molecular mechanisms and biological cues that govern the handling of their mRNA cargos. First, what developmental signaling initiates the IFE-3 to IFE-1 hand off that apparently precedes germ cell mRNA translational recruitment? Is eIF4E to eIF4E shuffling ever reversed? Does IFE-3 play an active role in miRNA placement or licensing of mRNAs transiting through germ granules? Since both IFE-1 and IFE-3 have single cap-binding ligand sites and very little RNA-binding capacity (87,112) and different 4EIP partners (40,46,48), how is the mRNA transfer accomplished? Is there mRNP remodeling in the germ granule, such that PGL-1 may play a role in moving IFE-1 out toward IFG-1? Is there a role for the helicase GLH-1 (22)? These are questions relevant to germ cell translational control that can now be addressed in the *C. elegans* and other germ cell systems.

## Supporting information

Suppl Fig 1 bioinformatics

Suppl Fig 2 western blot

Suppl Fig 3 RNAi

Suppl Fig 4 ife-5

## Acknowledgements

We would like to thank Dr David Greenstein (Univ of Minnesota) for providing the fluorescent reporter strains and Dr Peter Boag (Monash Univ) for the antibody to IFET-1, information on unpublished data and helpful discussions on the mRNP functions in germline development. Some of the strains for this study were provided by the CGC, funded by NIH Office of Research Infrastructure Programs (P40 OD010440). Image collection, processing and analysis for this manuscript was performed with the assistance of Dr. Frederic Bonnet and the MDI Biological Laboratory Light Microscopy Facility (LMF; RRID:SCR_019166).

## Data Availability Statement

Strains are available upon request. The authors affirm that all data necessary for confirming the conclusions of the article are present within the article, figures, and tables. The RNA-Seq and Mass Spectrometry data will be deposited at the GEO (https://www.ncbi.nlm.nih.gov/geo/) under the accession numbers are GSExxxxxx.

---

Supplemental Figure 1. <u>General bioinformatic analysis of IFE-1 vs IFE-3 mRNP RIP-Seq data</u>

A and B. Broader analysis of all represented transcripts with >2-fold enrichment and p<0.01 in RIP-Seq (right quadrant, Fig. 4B and C, purple dots) are further identified here for IFE-1 (A) and IFE-3 (B). The majority of sequences enriched in eIF4E mRNPs were highly abundant housekeeping mRNAs transcripts (red or brown dots, named) or unassigned genes in WormBase (purple dots). A. Ribosomal proteins L36A, L29, L34, S25, S17 mRNAs were 2-3-fold enriched in the IFE-1 mRNP with high significance (brown, p<10^−10^), as were a few with functions non-specific to the germline (red, *dct-16, nlp-24, hpo-18*). B. Histone mRNAs were identified as the predominant mRNAs enriched 2-5-fold in the IFE-3 mRNP (red dots, *his-1, −3, - 4, −31, −32, −48, −64* and *-67*, p<10^−10^), along with ribosomal protein L36A and 60S subunit acidic ribosomal protein (brown or off-scale, *rpl-36, rla-0*). Excluded mRNAs (left quadrant) were also noted. Among those excluded by both IFE-1 (A) and IFE-3 (B) are mRNAs encoding translation factors eEF1alpha (*eef-1A.1*), eEF2 (*eef-2*), the re-initiation factor ABCE-1 (*abce-1*), and IFE-3 itself (*ife-3*), as well as structural 5S rRNA (*rrn-4.1-11*). The other four *ife* mRNAs were not detected, and eIF4A (*inf-1*) was excluded from IFE-1 and not detected in IFE-3 pulldowns. This suggests that neither eIF4E exerts substantial autoregulatory translational repression feedback. Other general mRNAs including heat shock protein (*hsp-1*) and the other major tubulin (*tbb-1*) were excluded from both mRNPs. GO analysis comparing IFE mRNP to total mRNA identifies sequestered mRNAs clusters (C, D) that support a gene ratio analysis (E, F) to indicate “peptide metabolic” and “amide biosynthetic” processes (protein synthesis), cell cycle, reproduction, germ cell development, gamete generation, oogenesis, and mitotic processes. Separate gene ratio analysis for mRNAs direct comparison of IFE-3 over IFE-1 (G) focuses on reproduction, cell cycle, oogenesis, negative regulation of translation and embryonic pattern specification. H. Principle Components indicate a nominal clustering of the RNA-sequenced populations by sample type.

Supplemental <u>Figure 2. Efficient isolation of mKate2::IFE-1 and GFP::IFE-3 using affinity purification beads for RFP and GFP</u>

Western blots using anti-GFP and anti-RFP antibodies to detect isolation of the eIF4E fusion proteins using <u>Chromote</u>k immunoaffinity beads for sample preparation and precipitation for LC-MS proteomic analysis. Detection with HRP-conjugated secondary antibody used ECL+ followed by Typhoon Fluorescence/Phosphorimaging scanning. Western blotting for sample preparation and IP pulldown for RIP-Seq were similar (data not shown). The protein bound to 10% fraction of beads was compared to 20 ug aliquots of total protein for initial lysate and unbound fraction (approximately 1% by volume). Migration just above 50 kDa is consistent with the predicted relative masses of mKate2::3xMyc::IFE-1 and GFP::3xFlag::IFE-3 proteins.

Supplemental Figure 3. <u>Efficient depletion of PGL-1 by *pgl-1/3*(RNAi) without loss of IFE-1</u>

A. Depletion of PGL-1::GFP by the same *pgl-1/3*(RNAi) fed construct was very efficient at 25°C; Graphs of GFP quantification show 80-90% loss of PGL-1::GFP fluorescence in the distal gonad, proximal gonad and the embryo. Expression of mKate2::IFE-1 was unaffected by *pgl-1/3*(RNAi) at 25°C. Graphs of quantification of red mKate2::IFE-1 fluorescence showed insignificant signal decrease in the distal gonad, proximal gonad and the embryo upon *pgl-1/3*(RNAi).

Supplemental Figure 4. <u>Compensatory expression of the closely related *ife-5* gene does not occur in mutant backgrounds that deplete IFE-1 or IFE-3</u>

Total RNA extracted from worms treated with *ife-3*(RNAi), *ifet-1*(RNAi), *pgl-1/3*(RNAi) or bearing the *ife-1*(bn127) null mutation (as in Fig. 5) were subjected to qRT-PCR using primers specific for *ife-5* mRNA and *tbb-2* mRNA. No compensatory increase in *ife-5* gene expression was detected after loss of either IFE-1 or depletion of IFE-3 or IFET-1. There was a modest increase in *ife-5* mRNA after PGL-1/3 depletion. The nature of this almost two-fold increase is not clear.

## References

1. Mercer, M., Jang, S., Ni, C. and Buszczak, M. (2021) The Dynamic Regulation of mRNA Translation and Ribosome Biogenesis During Germ Cell Development and Reproductive Aging. Front Cell Dev Biol, 9, 710186.

2. Pelletier, J. and Sonenberg, N. (2019) The Organizing Principles of Eukaryotic Ribosome Recruitment. Annu. Rev. Biochem., 88, 307–335.

3. Keiper, B. (2019) Cap-Independent mRNA Translation in Germ Cells. Int J Mol Sci., 20, 173.

4. Browning, K.S. and Bailey-Serres, J. (2015) Mechanism of cytoplasmic mRNA translation. Arabidopsis Book, 13, e0176.

5. Keiper, B.D., Gan, W. and Rhoads, R.E. (1999) Protein synthesis initiation factor 4G. Int. J. Biochem. Cell Biol., 31, 37–41.

6. Gingras, A.-C., Raught, B. and Sonenberg, N. (1999) eIF4 initiation factors: Effectors of mRNA recruitment to ribosomes and regulators of translation. Annu. Rev. Biochem., 68, 913–963.

7. Hentze, M.W. (1997) eIF4G: a multipurpose ribosome adaptor? Science, 275, 500–501.

8. Merrick, W.C. and Hershey, J.W.B. (eds.) (1996) The pathway and mechanism of eukaryotic protein synthesis. Cold Spring Harbor Laboratory Press, Cold Spring Harbor.

9. Hellen, C.U.T. (2018) Translation Termination and Ribosome Recycling in Eukaryotes. Cold Spring Harb. Perspect. Biol., 10, a032656.

10. Mancera-Martinez, E., Brito Querido, J., Valasek, L.S., Simonetti, A. and Hashem, Y. (2017), RNA Biology. Taylor & Francis, pp. 1–7.

11. Richter, J.D. and Lasko, P. (2011) Translational control in oocyte development. Cold Spring Harb Perspect Biol, 3, a002758.

12. Huggins, H.P. and Keiper, B.D. (2020) Regulation of Germ Cell mRNPs by eIF4E:4EIP Complexes: Multiple Mechanisms, One Goal. Front Cell Dev Biol, 8, 562.

13. Sriram, A., Bohlen, J. and Teleman, A.A. (2018) Translation acrobatics: how cancer cells exploit alternate modes of translational initiation. EMBO Rep., 19, 17.

14. Clemens, M.J. (2004) Targets and mechanisms for the regulation of translation in malignant transformation. Oncogene, 23, 3180–3188.

15. Meric, F. and Hunt, K.K. (2002) Translation initiation in cancer: a novel target for therapy. Mol. Cancer Ther., 1, 971–979.

16. Iacoangeli, A. and Tiedge, H. (2013) Translational control at the synapse: role of RNA regulators. Trends Biochem Sci, 38, 47–55.

17. DeGracia, D.J., Kumar, R., Owen, C.R., Krause, G.S. and White, B.C. (2002) Molecular pathways of protein synthesis inhibition during brain reperfusion: implications for neuronal survival or death. J Cereb Blood Flow Metab, 22, 127–141.

18. Gkogkas, C.G., Khoutorsky, A., Ran, I., Rampakakis, E., Nevarko, T., Weatherill, D.B., Vasuta, C., Yee, S., Truitt, M., Dallaire, P. et al. (2013) Autism-related deficits via dysregulated eIF4E-dependent translational control. Nature, 493, 371–377.

19. Peter, D., Weber, R., Kane, C., Chung, M.Y., Ebertsch, L., Truffault, V., Weichenrieder, O., Igreja, C. and Izaurralde, E. (2015) Mextli proteins use both canonical bipartite and novel tripartite binding modes to form eIF4E complexes that display differential sensitivity to 4E-BP regulation. Genes Dev, 29, 1835–1849.

20. Chiluiza, D., Bargo, S., Callahan, R. and Rhoads, R.E. (2011) Expression of truncated eIF3e resulting from integration of MMTV causes a shift from Cap-dependent to Cap-independent translation. J. Biol. Chem., 286, 31288–31296.

21. Mayberry, L.K., Allen, M.L., Dennis, M.D. and Browning, K.S. (2009) Evidence for variation in the optimal translation initiation complex: plant eIF4B, eIF4F, and eIF(iso)4F differentially promote translation of mRNAs. Plant Physiol., 150, 1844–1854. doi: 1810.1104/pp.1109.138438. Epub 132009 Jun 138433.

22. Marnik, E.A., Fuqua, J.H., Sharp, C.S., Rochester, J.D., Xu, E.L., Holbrook, S.E. and Updike, D.L. (2019) Germline Maintenance Through the Multifaceted Activities of GLH/Vasa in Caenorhabditis elegans P Granules. Genetics, 213, 923–939. doi: 910.1534/genetics.1119.302670. Epub 302019 Sep 302610.

23. Mayya, V.K., Flamand, M.N., Lambert, A.M., Jafarnejad, S.M., Wohlschlegel, J.A., Sonenberg, N. and Duchaine, T.F. (2021) microRNA-mediated translation repression through GYF-1 and IFE-4 in C. elegans development. Nucleic Acids Res., gkab162.

24. Friday, A.J. and Keiper, B.D. (2015) Positive mRNA Translational Control in Germ Cells by Initiation Factor Selectivity. Biomed Research International, 2015, e327963.

25. Patrick, R.M., Mayberry, L.K., Choy, G., Woodard, L.E., Liu, J.S., White, A., Mullen, R.A., Tanavin, T.M., Latz, C.A. and Browning, K.S. (2014) Two Arabidopsis loci encode novel eukaryotic initiation factor 4E isoforms that are functionally distinct from the conserved plant eukaryotic initiation factor 4E. Plant Physiol., 164, 1820–1830.

26. Tettweiler, G., Kowanda, M., Lasko, P., Sonenberg, N. and Hernández, G. (2012) The Distribution of eIF4E-Family Members across Insecta. Comparative and Functional Genomics, 2012, 1–15.

27. Hernandez, G., Proud, C.G., Preiss, T. and Parsyan, A. (2012) On the Diversification of the Translation Apparatus across Eukaryotes. Comp. Funct. Genomics, 2012, 256848.

28. Hernandez, G., Altmann, M., Sierra, J.M., Urlaub, H., del Corral, R.D., Schwartz, P. and Rivera-Pomar, R. (2005) Functional analysis of seven genes encoding eight translation initiation factor 4E (eIF4E) isoforms in Drosophila. Mech. Dev., 122, 529–543.

29. Keiper, B.D., Lamphear, B.J., Deshpande, A.M., Jankowska-Anyszka, M., Aamodt, E.J., Blumenthal, T. and Rhoads, R.E. (2000) Functional characterization of five eIF4E isoforms in Caenorhabditis elegans. J. Biol. Chem., 275, 10590–10596.

30. Nousch, M. and Eckmann, C.R. (2013) Translational control in the Caenorhabditis elegans germ line. Adv. Exp. Med. Biol., 757, 205–247.

31. Gao, M. and Arkov, A.L. (2013) Next generation organelles: structure and role of germ granules in the germline. Mol Reprod Dev, 80, 610–623. doi: 610.1002/mrd.22115. Epub 22012 Nov 22113.

32. Marnik, E.A. and Updike, D.L. (2019) Membraneless organelles: P granules in Caenorhabditis elegans. Traffic, 20, 373–379.

33. Phillips, C.M. and Updike, D.L. (2022) Germ granules and gene regulation in the Caenorhabditis elegans germline. Genetics, 220, iyab195.

34. Uebel, C.J., Rajeev, S. and Phillips, C.M. (2023) Caenorhabditis elegans germ granules are present in distinct configurations and assemble in a hierarchical manner. Development, 150.

35. Uebel, C.J., Manage, K.I. and Phillips, C.M. (2021) SIMR foci are found in the progenitor germ cells of C. elegans embryos. MicroPubl Biol, 2021.

36. Wan, G., Fields, B.D., Spracklin, G., Shukla, A., Phillips, C.M. and Kennedy, S. (2018) Spatiotemporal regulation of liquid-like condensates in epigenetic inheritance. Nature, 557, 679–683.

37. Lee, C.S., Putnam, A., Lu, T., He, S., Ouyang, J.P.T. and Seydoux, G. (2020) Recruitment of mRNAs to P granules by condensation with intrinsically-disordered proteins. Elife, 9, e52896.

38. Brangwynne, C.P., Eckmann, C.R., Courson, D.S., Rybarska, A., Hoege, C., Gharakhani, J., Julicher, F. and Hyman, A.A. (2009) Germline P granules are liquid droplets that localize by controlled dissolution/condensation. Science., 324, 1729–1732. doi: 1710.1126/science.1172046. Epub 1172009 May 1172021.

39. Pushpa, K., Kumar, G.A. and Subramaniam, K. (2017) Translational Control of Germ Cell Decisions. Results Probl. Cell Differ., 59:175–200., 10.1007/1978-1003-1319-44820-44826_44826.

40. Huggins, H.P., Subash, J.S., Stoffel, H., Henderson, M.A., Hoffman, J.L., Buckner, D.S., Sengupta, M.S., Boag, P.R., Lee, M.H. and Keiper, B.D. (2020) Distinct roles of two eIF4E isoforms in the germline of Caenorhabditis elegans. J. Cell. Sci., 133(6). jcs.237990. doi: 237910.231242/jcs.237990.

41. Friday, A.J., Henderson, M.A., Morrison, J.K., Hoffman, J.L. and Keiper, B.D. (2015) Spatial and temporal translational control of germ cell mRNAs mediated by the eIF4E isoform IFE-1. J. Cell Sci., 128, 4487–4498.

42. Song, A., Labella, S., Korneeva, N.L., Keiper, B.D., Aamodt, E.J., Zetka, M. and Rhoads, R.E. (2010) A C. elegans eIF4E-family member upregulates translation at elevated temperatures of mRNAs encoding MSH-5 and other meiotic crossover proteins. J. Cell Sci., 123, 2228–2237.

43. Dinkova, T.D., Keiper, B.D., Korneeva, N.L., Aamodt, E.J. and Rhoads, R.E. (2005) Translation of a small subset of Caenorhabditis elegans mRNAs is dependent on a specific eukaryotic translation initiation factor 4E isoform. Mol. Cell. Biol., 25, 100–113.

44. Henderson, M.A., Cronland, E., Dunkelbarger, S., Contreras, V., Strome, S. and Keiper, B.D. (2009) A germ line-specific isoform of eIF4E (IFE-1) is required for efficient translation of stored mRNAs and maturation of both oocytes and sperm. J. Cell Science, 122, 1529–1539.

45. Marnik, E.A., Almeida, M.V., Cipriani, P.G., Chung, G., Caspani, E., Karaulanov, E., Gan, H.H., Zinno, J., Isolehto, I.J., Kielisch, F. et al. (2022) The Caenorhabditis elegans TDRD5/7-like protein, LOTR-1, interacts with the helicase ZNFX-1 to balance epigenetic signals in the germline. PLoS Genet, 18, e1010245.

46. Amiri, A., Keiper, B.D., Kawasaki, I., Fan, Y., Kohara, Y., Rhoads, R.E. and Strome, S. (2001) An isoform of eIF4E is a component of germ granules and is required for spermatogenesis in *C. elegans*. Development, 128, 3899–3912.

47. Brenner, S. (1974) The genetics of Caenorhabditis elegans. Genetics, 77, 71–94.

48. Sengupta, M.S., Low, W.Y., Patterson, J.R., Kim, H.M., Traven, A., Beilharz, T.H., Colaiacovo, M.P., Schisa, J.A. and Boag, P.R. (2013) ifet-1 is a broad-scale translational repressor required for normal P granule formation in C. elegans. J. Cell Sci., 126, 850–859.

49. Sengupta, M.S. and Boag, P.R. (2012) Germ granules and the control of mRNA translation. IUBMB Life, 64, 586–594.

50. Hanazawa, M., Kawasaki, I., Kunitomo, H., Gengyo-Ando, K., Bennett, K.L., Mitani, S. and Iino, Y. (2004) The Caenorhabditis elegans eukaryotic initiation factor 5A homologue, IFF-1, is required for germ cell proliferation, gametogenesis and localization of the P-granule component PGL-1. Mech Dev, 121, 213–224.

51. Kawasaki, I., Amiri, A., Fan, Y., Meyer, N., Dunkelbarger, S., Motohashi, T., Karashima, T., Bossinger, O. and Strome, S. (2004) The PGL family proteins associate with germ granules and function redundantly in Caenorhabditis elegans germline development. Genetics, 167, 645–661.

52. Updike, D.L., Knutson, A.K., Egelhofer, T.A., Campbell, A.C. and Strome, S. (2014) Germ-granule components prevent somatic development in the C. elegans germline. Curr. Biol., 24, 970–975.

53. Hickey, K.L., Dickson, K., Cogan, J.Z., Replogle, J.M., Schoof, M., D’Orazio, K.N., Sinha, N.K., Hussmann, J.A., Jost, M., Frost, A. et al. (2020) GIGYF2 and 4EHP Inhibit Translation Initiation of Defective Messenger RNAs to Assist Ribosome-Associated Quality Control. Mol Cell, 79, 950–962 e956.

54. Hernandez, G., Miron, M., Han, H., Liu, N., Magescas, J., Tettweiler, G., Frank, F., Siddiqui, N., Sonenberg, N. and Lasko, P. (2013) Mextli is a novel eukaryotic translation initiation factor 4E-binding protein that promotes translation in Drosophila melanogaster. Mol. Cell. Biol., 33, 2854–2864.

55. Contreras, V., Richardson, M.A., Hao, E. and Keiper, B.D. (2008) Depletion of the cap-associated isoform of translation factor eIF4G induces germline apoptosis in C. elegans. Cell Death Differ., 15, 1232–1242.

56. Morrison, J.K., Friday, A.J., Henderson, M.A., Hao, E. and Keiper, B.D. (2014) Induction of cap-independent BiP (*hsp-3*) and Bcl-2 (*ced-9*) translation in response to eIF4G (IFG-1) depletion in *C. elegans*. Translation, 2, e28935.

57. Li, Y., Snyder, M. and Maine, E.M. (2021) Meiotic H3K9me2 distribution is influenced by the ALG-3 and ALG-4 pathway and by poly(U) polymerase activity. MicroPubl Biol, 2021.

58. van Wolfswinkel, J.C., Claycomb, J.M., Batista, P.J., Mello, C.C., Berezikov, E. and Ketting, R.F. (2009) CDE-1 affects chromosome segregation through uridylation of CSR-1-bound siRNAs. Cell, 139, 135–148.

59. Wang, Y., Weng, C., Chen, X., Zhou, X., Huang, X., Yan, Y. and Zhu, C. (2020) CDE-1 suppresses the production of risiRNA by coupling polyuridylation and degradation of rRNA. BMC Biol, 18, 115.

60. Zheng, H., Peng, K., Gou, X., Ju, C. and Zhang, H. (2023) RNA recruitment switches the fate of protein condensates from autophagic degradation to accumulation. J Cell Biol, 222.

61. Cordeiro Rodrigues, R.J., de Jesus Domingues, A.M., Hellmann, S., Dietz, S., de Albuquerque, B.F.M., Renz, C., Ulrich, H.D., Sarkies, P., Butter, F. and Ketting, R.F. (2019) PETISCO is a novel protein complex required for 21U RNA biogenesis and embryonic viability. Genes Dev., 33, 857–870.

62. Perez-Borrajero, C., Podvalnaya, N., Holleis, K., Lichtenberger, R., Karaulanov, E., Simon, B., Basquin, J., Hennig, J., Ketting, R.F. and Falk, S. (2021) Structural basis of PETISCO complex assembly during piRNA biogenesis in C. elegans. Genes Dev, 35, 1304–1323.

63. Gray, N.K. and Wickens, M. (1998) Control of translation initiation in animals. Annu Rev Cell Dev Biol, 14, 399–458.

64. Cauchi, R.J. (2010) SMN and Gemins:‘we are family’… or are we? Insights into the partnership between Gemins and the spinal muscular atrophy disease protein SMN. Bioessays, 32, 1077–1089.

65. Tsukamoto, T., Gearhart, M.D., Spike, C.A., Huelgas-Morales, G., Mews, M., Boag, P.R., Beilharz, T.H. and Greenstein, D. (2017) LIN-41 and OMA Ribonucleoprotein Complexes Mediate a Translational Repression-to-Activation Switch Controlling Oocyte Meiotic Maturation and the Oocyte-to-Embryo Transition in Caenorhabditis elegans. Genetics, 206, 2007–2039.

66. Ogura, K., Kishimoto, N., Mitani, S., Gengyo-Ando, K. and Kohara, Y. (2003) Translational control of maternal glp-1 mRNA by POS-1 and its interacting protein SPN-4 in Caenorhabditis elegans. Development, 130, 2495–2503.

67. Albarqi, M.M.Y. and Ryder, S.P. (2021) The endogenous mex-3 3’UTR is required for germline repression and contributes to optimal fecundity in C. elegans. PLoS Genet, 17, e1009775.

68. Albarqi, M.M.Y. and Ryder, S.P. (2023) The role of RNA-binding proteins in orchestrating germline development in Caenorhabditis elegans. Front Cell Dev Biol, 10, 1094295.

69. Kawasaki, I., Shim, Y.H., Kirchner, J., Kaminker, J., Wood, W.B. and Strome, S. (1998) PGL-1, a predicted RNA-binding component of germ granules, is essential for fertility in *C. elegans*. Cell, 94, 635–645.

70. Kawasaki, I., Jeong, M.H. and Shim, Y.H. (2011) Regulation of sperm-specific proteins by IFE-1, a germline-specific homolog of eIF4E, in C. elegans. Mol. Cells, 31, 191–197.

71. Navarro, R.E., Shim, E.Y., Kohara, Y., Singson, A. and Blackwell, T.K. (2001) cgh-1, a conserved predicted RNA helicase required for gametogenesis and protection from physiological germline apoptosis in C. elegans. Development, 128, 3221–3232.

72. Mendez, R. and Richter, J.D. (2001) Translational control by CPEB: a means to the end. Nat. Rev. Mol. Cell Biol., 2, 521–529.

73. Stebbins-Boaz, B., Cao, Q., de Moor, C.H., Mendez, R. and Richter, J.D. (1999) Maskin is a CPEB-associated factor that transiently interacts with elF-4E. Mol. Cell, 4, 1017–1027.

74. Keiper, B.D. and Rhoads, R.E. (1999) Translational recruitment of *Xenopus* maternal mRNAs in response to poly(A) elongation requires initiation factor eIF4G-1. Dev. Biol., 206, 1–14.

75. Prevot, D., Darlix, J.L. and Ohlmann, T. (2003) Conducting the initiation of protein synthesis: the role of eIF4G. Biol. Cell, 95, 141–156.

76. Cao, Q. and Richter, J.D. (2002) Dissolution of the maskin-eIF4E complex by cytoplasmic polyadenylation and poly(A)-binding protein controls cyclin B1 mRNA translation and oocyte maturation. EMBO J., 21, 3852–3862.

77. Tabara, H., Hill, R.J., Mello, C.C., Priess, J.R. and Kohara, Y. (1999) pos-1 encodes a cytoplasmic zinc-finger protein essential for germline specification in C. elegans. Development, 126, 1–11.

78. Draper, B.W., Mello, C.C., Bowerman, B., Hardin, J. and Priess, J.R. (1996) MEX-3 is a KH domain protein that regulates blastomere identity in early C. elegans embryos. Cell, 87, 205–216.

79. Shimada, M., Yokosawa, H. and Kawahara, H. (2006) OMA-1 is a P granules-associated protein that is required for germline specification in Caenorhabditis elegans embryos. Genes Cells, 11, 383–396.

80. Schisa, J.A., Pitt, J.N. and Priess, J.R. (2001) Analysis of RNA associated with P granules in germ cells of C. elegans adults. Development, 128, 1287–1298.

81. Spike, C.A., Coetzee, D., Eichten, C., Wang, X., Hansen, D. and Greenstein, D.I. (2014) The TRIM-NHL Protein LIN-41 and the OMA RNA-Binding Proteins Antagonistically Control the Prophase-to-Metaphase Transition and Growth of Caenorhabditis elegans Oocytes. Genetics, 198, 1535–1558.

82. Tocchini, C., Keusch, J.J., Miller, S.B., Finger, S., Gut, H., Stadler, M.B. and Ciosk, R. (2014) The TRIM-NHL protein LIN-41 controls the onset of developmental plasticity in Caenorhabditis elegans. PLoS Genet, 10, e1004533 (epub ahead of print).

83. Merrick, W.C. (1992) Mechanism and regulation of eukaryotic protein synthesis. Microbiol. Reviews, 56, 291–315.

84. Gruner, S., Peter, D., Weber, R., Wohlbold, L., Chung, M.Y., Weichenrieder, O., Valkov, E., Igreja, C. and Izaurralde, E. (2016) The Structures of eIF4E-eIF4G Complexes Reveal an Extended Interface to Regulate Translation Initiation. Mol. Cell, 2765, 020.

85. Aoki, S.T., Lynch, T.R., Crittenden, S.L., Bingman, C.A., Wickens, M. and Kimble, J. (2021) C. elegans germ granules require both assembly and localized regulators for mRNA repression. Nature communications, 12, 996.

86. Frydryskova, K., Masek, T., Borcin, K., Mrvova, S., Venturi, V. and Pospisek, M. (2016) Distinct recruitment of human eIF4E isoforms to processing bodies and stress granules. BMC Mol. Biol., 17, 21.

87. Miyoshi, H., Dwyer, D.S., Keiper, B.D., Jankowska-Anyszka, M., Darzynkiewicz, E. and Rhoads, R.E. (2002) Discrimination between mono- and trimethylated cap structures by two isoforms of *Caenorhabditis elegans* eIF4E. EMBO J., 21, 4680–4690.

88. Patrick, R.M. and Browning, K.S. (2012) The eIF4F and eIFiso4F Complexes of Plants: An Evolutionary Perspective. Comp. Funct. Genomics, 2012, 287814.

89. Contreras, V., Friday, A.J., Morrison, J.K., Hao, E. and Keiper, B.D. (2011) Cap-Independent translation promotes *C. elegans* germ cell apoptosis through Apaf-1/CED-4 in a caspase-dependent mechanism. PLoS ONE, 6, e24,444.

90. Dai, S., Tang, X., Li, L., Ishidate, T., Ozturk, A.R., Chen, H., Dube, A.L., Yan, Y.-H., Dong, M.-Q. and Shen, E.-Z. (2022) A family of C. elegans VASA homologs control Argonaute pathway specificity and promote transgenerational silencing. Cell reports, 40.

91. Spichal, M., Heestand, B., Billmyre, K.K., Frenk, S., Mello, C.C. and Ahmed, S. (2021) Germ granule dysfunction is a hallmark and mirror of Piwi mutant sterility. Nature Communications, 12, 1420.

92. Li, W., DeBella, L.R., Guven-Ozkan, T., Lin, R. and Rose, L.S. (2009) An eIF4E-binding protein regulates katanin protein levels in C. elegans embryos. J Cell Biol, 187, 33–42.

93. Mangio, R.S., Votra, S. and Pruyne, D. (2015) The canonical eIF4E isoform of C. elegans regulates growth, embryogenesis, and germline sex-determination. Biol. Open, 4, 843–851.

94. Goh, W.-S.S., Seah, J.W.E., Harrison, E.J., Chen, C., Hammell, C.M. and Hannon, G.J. (2014) A genome-wide RNAi screen identifies factors required for distinct stages of C. elegans piRNA biogenesis. Genes & development, 28, 797–807.

95. Jankowska-Anyszka, M., Lamphear, B.J., Aamodt, E.J., Harrington, T., Darzynkiewicz, E., Stolarski, R. and Rhoads, R.E. (1998) Multiple isoforms of eukaryotic protein synthesis initiation factor 4E in *Caenorhabditis elegans* can distinguish between, mono- and trimethylated mRNA cap structures. J. Biol. Chem., 273, 10538–10542.

96. Thoreen, C.C., Chantranupong, L., Keys, H.R., Wang, T., Gray, N.S. and Sabatini, D.M. (2012) A unifying model for mTORC1-mediated regulation of mRNA translation. Nature, 485, 109–113.

97. Roy, D., Kahler, D.J., Yun, C. and Hubbard, E.J.A. (2018) Functional Interactions Between rsks-1/S6K, glp-1/Notch, and Regulators of Caenorhabditis elegans Fertility and Germline Stem Cell Maintenance. G3 (Bethesda), 8, 3293–3309.

98. Pan, K.Z., Palter, J.E., Rogers, A.N., Olsen, A., Chen, D., Lithgow, G.J. and Kapahi, P. (2007) Inhibition of mRNA translation extends lifespan in Caenorhabditis elegans. Aging Cell, 6, 111–119.

99. Barnard, D.C., Ryan, K., Manley, J.L. and Richter, J.D. (2004) Symplekin and xGLD-2 are required for CPEB-mediated cytoplasmic polyadenylation. Cell., 119, 641–651. doi: 610.1016/j.cell.2004.1010.1029.

100. Cui, J., Sartain, C.V., Pleiss, J.A. and Wolfner, M.F. (2013) Cytoplasmic polyadenylation is a major mRNA regulator during oogenesis and egg activation in Drosophila. Dev Biol, 383, 121–131. doi: 110.1016/j.ydbio.2013.1008.1013. Epub 2013 Aug 1023.

101. Kim, K.W., Wilson, T.L. and Kimble, J. (2010) GLD-2/RNP-8 cytoplasmic poly(A) polymerase is a broad-spectrum regulator of the oogenesis program. Proc Natl Acad Sci U S A, 107, 17445–17450. doi: 17410.11073/pnas.1012611107. Epub 1012612010 Sep 1012611120.

102. Nousch, M., Yeroslaviz, A. and Eckmann, C.R. (2019) Stage-specific combinations of opposing poly(A) modifying enzymes guide gene expression during early oogenesis. Nucleic Acids Res, 47, 10881–10893.

103. Nousch, M., Yeroslaviz, A., Habermann, B. and Eckmann, C.R. (2014) The cytoplasmic poly(A) polymerases GLD-2 and GLD-4 promote general gene expression via distinct mechanisms. Nucleic Acids Res., 42, 11622–11633.

104. Millonigg, S., Minasaki, R., Nousch, M., Novak, J. and Eckmann, C.R. (2014) GLD-4-mediated translational activation regulates the size of the proliferative germ cell pool in the adult C. elegans germ line. PLoS Genet, 10, e1004647.

105. Rybarska, A., Harterink, M., Jedamzik, B., Kupinski, A.P., Schmid, M. and Eckmann, C.R. (2009) GLS-1, a novel P granule component, modulates a network of conserved RNA regulators to influence germ cell fate decisions. PLoS Genet, 5, e1000494.

106. Davis, M.R., Delaleau, M. and Borden, K.L.B. (2019) Nuclear eIF4E Stimulates 3’-End Cleavage of Target RNAs. Cell Rep, 27, 1397–1408.e1394.

107. Zinshteyn, B., Sinha, N.K., Enam, S.U., Koleske, B. and Green, R. (2021) Translational repression of NMD targets by GIGYF2 and EIF4E2. PLoS Genet, 17, e1009813.

108. Christie, M. and Igreja, C. (2023) eIF4E-homologous protein (4EHP): a multifarious cap-binding protein. The FEBS Journal, 290, 266–285.

109. Long, X., Spycher, C., Han, Z.S., Rose, A.M., Muller, F. and Avruch, J. (2002) TOR deficiency in C. elegans causes developmental arrest and intestinal atrophy by inhibition of mRNA translation. Curr. Biol., 12, 1448–1461.

110. Flamand, M.N., Wu, E., Vashisht, A., Jannot, G., Keiper, B.D., Simard, M.J., Wohlschlegel, J. and Duchaine, T.F. (2016) Poly(A)-binding proteins are required for microRNA-mediated silencing and to promote target deadenylation in C. elegans. Nucleic Acids Res, 44, 5924–5935.

111. Rogers, A.N., Chen, D., McColl, G., Czerwieniec, G., Felkey, K., Gibson, B.W., Hubbard, A., Melov, S., Lithgow, G.J. and Kapahi, P. (2011) Life span extension via eIF4G inhibition is mediated by posttranscriptional remodeling of stress response gene expression in C. elegans. Cell Metab, 14, 55–66.

112. Marcotrigiano, J., Gingras, A.-C., Sonenberg, N. and Burley, S.K. (1997) Cocrystal structure of the messenger RNA 5’ cap-binding protein (eIF4E) bound to 7-methyl-GDP. Cell, 89, 951–961.

