## Supplementary figures and images for "Two germ granule eIF4E isoforms reside in different mRNPs to hand off *C elegans* mRNAs from translational repression to activation"

### Suppl Fig 1 bioinformatics

#### Supplement Figure 1

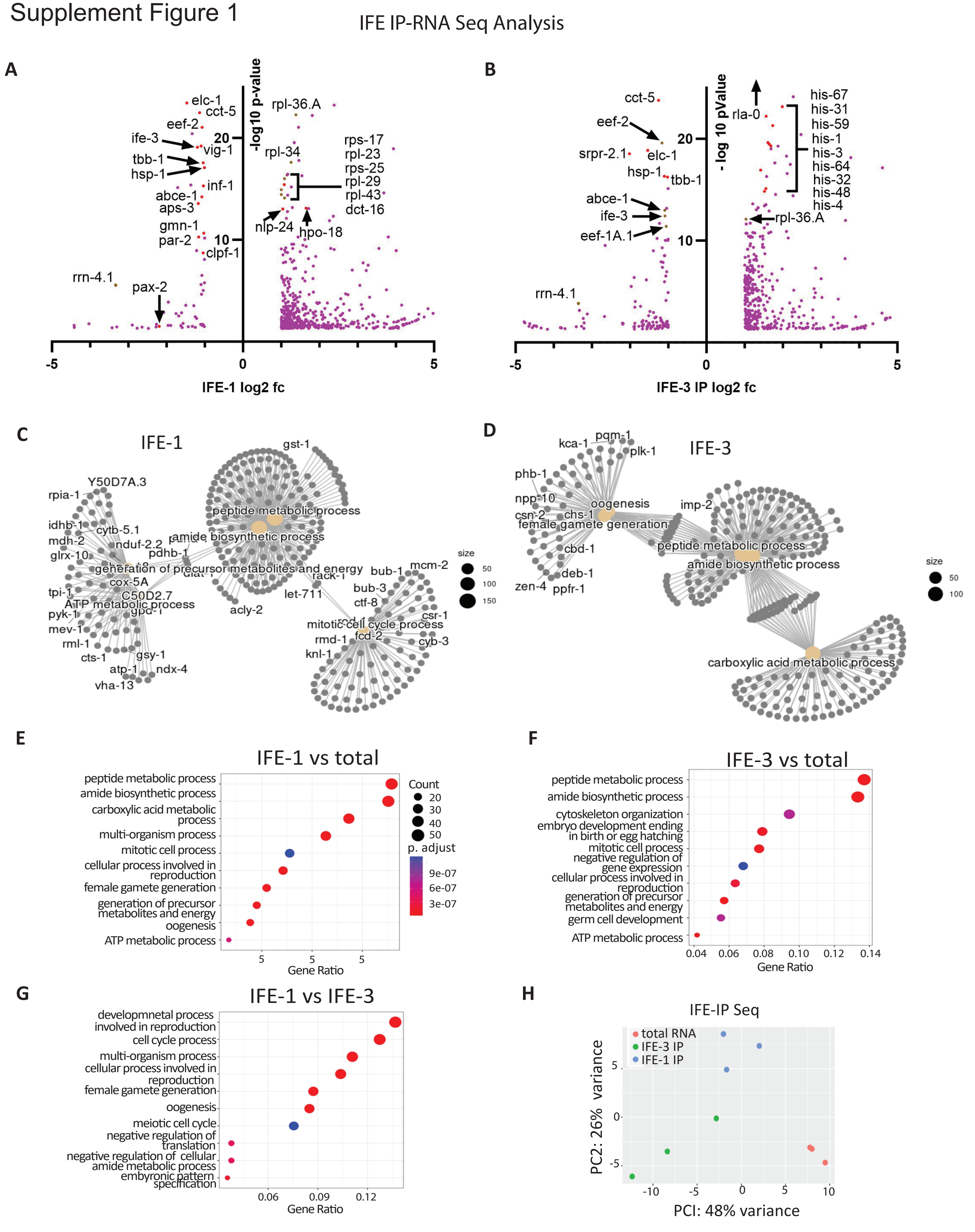

### Suppl Fig 2 western blot

Supplemental Figure 2

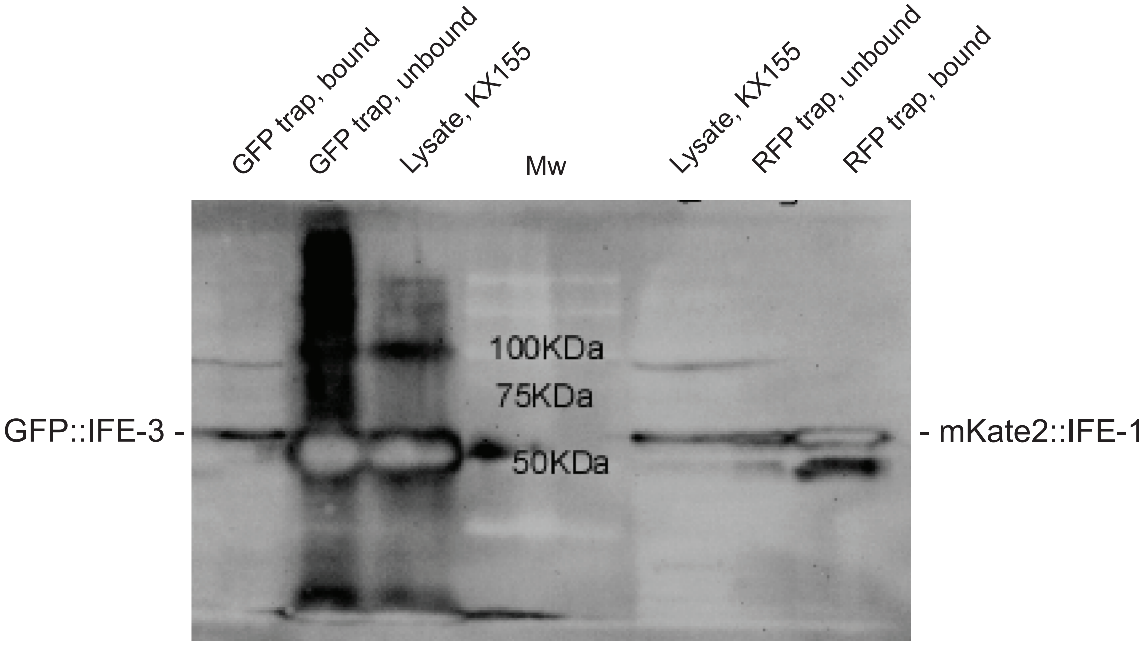

### Suppl Fig 3 RNAi

#### Supplemental Figure 3

##### Efficient PGL-1 depletion with no loss of IFE-1 at 25C

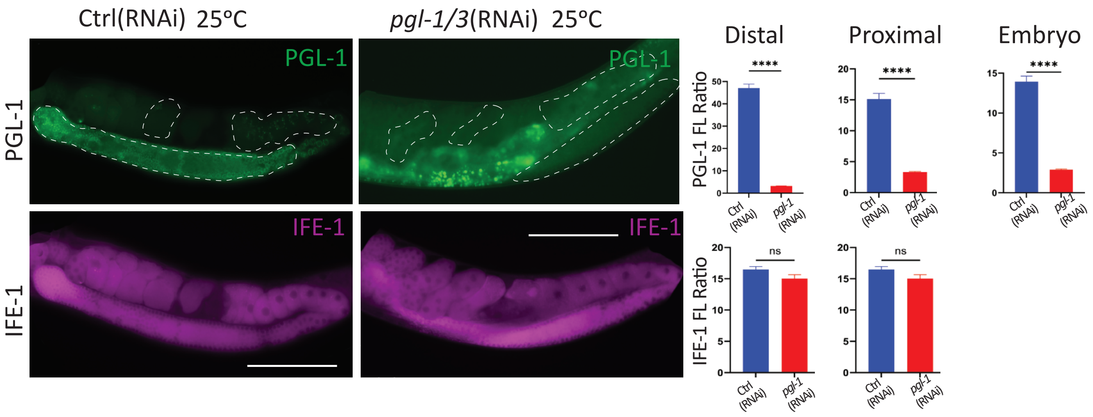

### Suppl Fig 4 ife-5

Supplement Figure 4

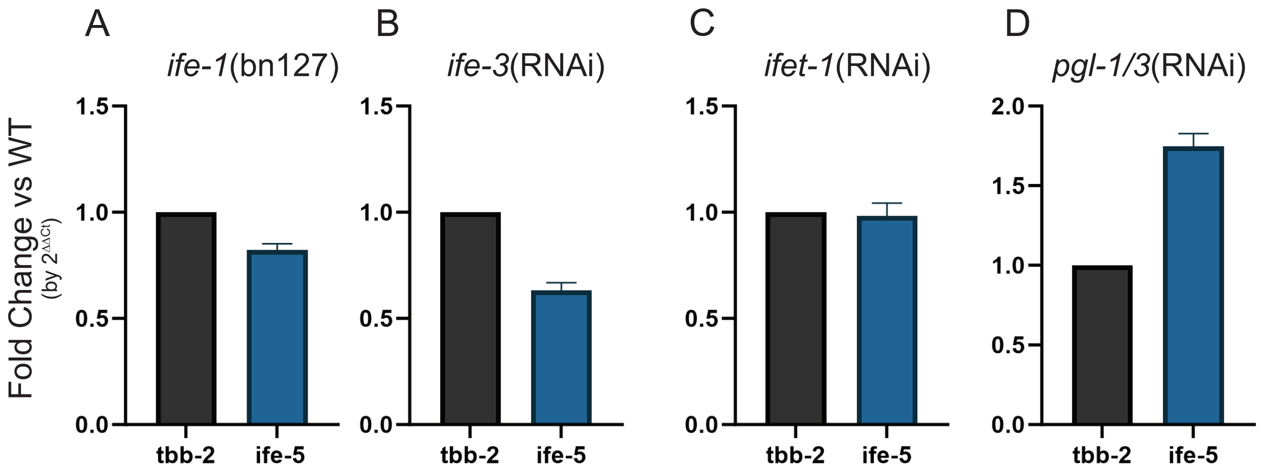
